# A cAMP/PKA-dependent synaptically targeted lncRNA mediates structural plasticity in hippocampal neurons by functionally interacting with the Spectrin/Ankyrin Network

**DOI:** 10.1101/2020.09.10.291526

**Authors:** Eddie Grinman, Yoshihisa Nakahata, Yosef Avchalumov, Isabel Espadas, Supriya Swarnkar, Ryohei Yasuda, Sathyanarayanan V. Puthanveettil

**Affiliations:** Department of Neuroscience, Scripps Research, 130 Scripps Way, Jupiter FL, 33458, USA; Max Planck Florida Institute for Neuroscience, Jupiter, FL 33458, USA

## Abstract

Activity-dependent structural plasticity at the synapse requires specific changes in the neuronal transcriptome. While much is known about the role of coding elements in this process, the role of the long-noncoding transcriptome remains elusive. Here we report the discovery of an intronic long noncoding RNA (lncRNA)—termed ADEPTR—whose expression is upregulated and is synaptically transported in a cAMP/PKA-dependent manner in hippocampal neurons, independent of its protein-coding host gene. Loss of ADEPTR function suppresses activity-dependent changes in synaptic transmission and structural plasticity of dendritic spines. Mechanistically, dendritic localization of ADEPTR is mediated by molecular motor protein Kif2A. ADEPTR physically binds to actin-scaffolding regulators Ankyrin (AnkB) and Spectrin (Sptn1) and is required for their dendritic localization. Taken together, this study demonstrates that ADEPTR regulates the dendritic Spectrin-Ankyrin network for structural plasticity at the synapse and illuminates a novel role for lncRNAs at the synapse.

**One Sentence Summary:** We have uncovered an intronic long noncoding RNA that is synaptically transported in a cAMP-dependent manner and is linked to cytoskeletal components of structural plasticity in hippocampal neurons.

## Introduction

Activity-dependent structural plasticity of neuronal synapses is a critical component of adaptive brain functions, such as long-term memory, and requires changes in the expression of transcriptional networks. The protein-coding factors of this transcriptional response to neuronal activation have been well studied, including Arc [1], Camk2a [2], Homer1a [3], BDNF [4] and more, that are distally transported and translated locally at activated synapses [5, 6]. These gene products then facilitate the structural reorganization therein by acting as signaling factors, scaffolds for the cytoskeleton, receptor subunits and more [7].

Recent advances in next generation sequencing studies have led to a deeper understanding of the components of the transcriptome, notably that it contains several thousands of large RNA products [8] that do not code for protein [9]. While the vast majority of these long-noncoding RNAs (lncRNAs) have yet to be experimentally interrogated and their role in mediating synapse function poorly understood, lncRNAs have previously been profiled that contribute to neuronal function by regulating target mRNA or rRNA transcription and processing in the nucleus [10, 11], or interacting with miRNAs in the cytoplasm [12]. Despite these advances, it remains unclear whether and how lncRNAs mediates activity dependent changes at the synapse.

A major consequence of activity dependent transcriptional changes is the transport of RNAs to synapses for remodeling of pre-existing synapses and formation of new ones. Though much is known about the significance of RNAs that are transported [13, 14] from the soma to the synapse for local translation and structural plasticity, it is unknown whether lncRNAs are transported to synapses for mediating local changes. Of intrigue, lncRNAs are known to display activity-dependent responses similar to mRNAs in neurons [15] but show higher turnover rates [16]. However, it is unknown whether and how lncRNAs can be distally transported during activity-dependent plasticity changes at the synapse.

Here we report the discovery as well as the functional and mechanistic characterization of an activity dependent lncRNA that is distally transported in mouse hippocampal neurons. We found that this lncRNA is rapidly expressed and transported to distal processes, is necessary for activity-dependent structural plasticity, and functionally interacts with the Spectrin/Ankyrin network in dendrites through a domain conserved in human 3’ UTRs.

## Results

### Discovery of an activity dependent synaptically targeted lncRNA

We employed a multipronged approach to begin uncovering lncRNAs that are robustly expressed and transported in activated hippocampal neurons, summarized in Figure 1A. We first profiled lncRNA expression in mature primary hippocampal neurons that were exposed to Forskolin to activate cAMP signaling, a signaling cascade known to mediate long term potentiation in cultured neurons and learning *in vivo* [17–19]. Total RNA isolated from three biological replicates per group was subjected to next generation RNA sequencing, clustering and differential expression analysis (Fig. S1A,B). In addition to previously known immediate early genes (Fos, Egr2, Nr4a1, and Arc), using gene biotype analysis we observed numerous Ensembl-annotated lncRNAs from varying genomic origins (Fig. S1B). We then asked whether cAMP regulated lncRNAs are enriched distally from the soma by employing a fractionation strategy to generate hippocampal synaptoneurosomes (Figs. S2A, B), isolate the RNA therein (Fig. S2C) and systematically screen lncRNA candidates by qRT-PCR (Fig. S2D, Supplementary Table S1). Analysis of synaptoneurosomes from neurons exposed to Forskolin show that Gm38257 was the most robustly expressed and consistently enriched in a cAMP-dependent manner [Fold change in Gm38257 expression in FSK synaptosomes compared to DMSO total lysate: 16.38 ± 2.26, p<0.001, DMSO synaptosomes compared to DMSO total lysate:1.55 ± 0.48, p = 0.98, Dunnett’s Test, N = 3-4 replicates per group] approaching fold enrichment levels similar to Arc mRNA (Fig. 1B, Supplementary Table S1).

**Figure 1:**
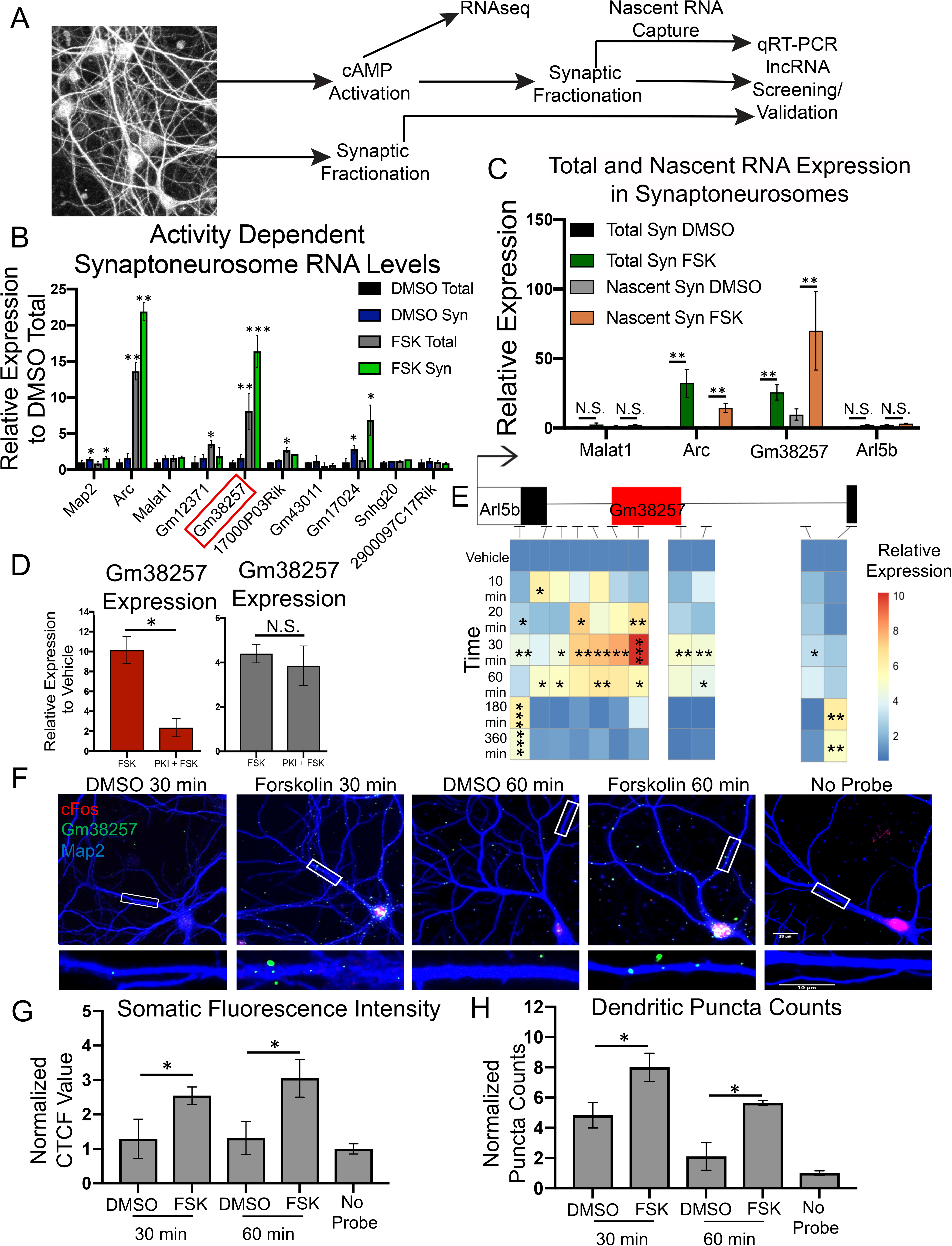
ADEPTR is a novel lncRNA that is expressed in hippocampal neurons and dendritically localized in a cAMP-dependent manner. (A) Schematic of multi-pronged approach to identify activity-dependent synaptically targeted lncRNAs. (B) Cell fractions or total lysates from FSK (50 uM) or DMSO 30 min treated neurons were collected in Trizol for RNA extraction. qRT-PCR was performed on N=3-4 replicates using relative quantification with 18s RNA serving as an internal control. ** p<0.01, * p<0.05, Dunnett’s Test compared to DMSO Total. (C) Primary hippocampal neurons were treated with EU-labeled nucleotides for incorporation into nascent transcripts followed by Forskolin or DMSO treatment and synaptosome preparation, as in (B). Total synaptic RNA was extracted prior to click-chemistry based nascent RNA capture. Malat1 and Arc were used as negative and positive controls, respectively. One Way-ANOVA, Tukey’s HSD, N=6-7 per group from 3 independent experiments. **(D)** DIV 16 primary hippocampal neurons were treated Forskolin (50 uM) for 30 min or 14-22 amide (PKI, 25uM) for 30 min followed by Forskolin (50uM) for another 30 min prior to total RNA isolation and qRT-PCR. * p <0.05, Unpaired Two-Sided T-Test. N=3-4 per group. **(E)** Heatmap showing expression of Arl5b exons and multiple introns—including ADEPTR—at the indicated time points, relative to vehicle. Primary hippocampal neurons DIV 16 were treated with Forskolin for the indicated times, followed by total RNA extraction and qRT-PCR analysis. Asterisks represent statistical significance relative to vehicle by Dunnett’s Test. N=3-5 per group. **(F)** Neurons were fixed and processed for FISH using digoxygenin labeled riboprobes. Cells were immunolabeled with Map2 to label dendrites and cFos to measure neuronal activation. (**G**) Somatic fluorescence intensity was measured using CTCF measurement in ImageJ. **(H)** Extra somatic puncta were acquired by a custom ImageJ macros script; counts were normalized to no-probe control. N=17-24 per group from 3 independent experiments. *p<0.05, **p<0.01, Pairwise T-test with Holm’s correction. Error bars throughout represent SEM. Scale bar represents 20 microns in the neuron image and 10 microns in the inset.

### Gm38257 is expressed and localized to synaptic compartments in a cAMP-dependent manner

Inspection of the genomic locus of Gm38257 revealed that it is a single exon transcript located within the first intron of Arl5b, a protein-coding gene (Fig. S3A). PhyloCSF [20] and CPAT [21] protein coding assessment tools both showed that Gm38257 is not likely to code for protein (Figs. S3A, B). qRT-PCR analysis from nuclear and cytoplasmic fractions revealed that both Gm38257 and Arl5b are upregulated by cAMP signaling and are expressed in both nuclear and cytoplasmic fractions (Fig. S4A). We next asked whether newly synthesized Gm38257 is distally targeted by nascent RNA capture analysis. Our measurements of total and nascent RNA captured from synaptoneurosomes, show that Gm38257, but not Arl5b, is enriched in synaptoneurosomes upon Forskolin stimulation [Supplementary Table S1, Gm38257 Nascent RNA with FSK vs Nascent RNA with DMSO: p = 0.02, One-Way ANOVA followed by Tukey’s HSD; Arl5b: p = 0.08 One Way ANOVA, N=6-7 per group] (Fig. 1C, Supplementary Table S1).

By performing a time course of cAMP-signaling in hippocampal neurons and designing qPCR primers tiling the exon 1, 2 and intron 1 of Arl5b, we found that Gm38257 and Arl5b have alternate expression patterns (Fig. 1D, Supplementary Table S1). We further determined that cAMP dependent Gm38257 expression, but not Arl5b expression, is reduced by inhibition of PKA with 14-22 amide [Gm38257 FSK+PKI vs Gm38257 FSK: p = 0.012, Arl5b FSK+PKI vs Arl5b FSK: p = 0.72, Two tailed unpaired T-test, N=3-4 per group] (Fig. 1E, Supplementary Table S1), suggesting that Gm38257 expression is regulated by cAMP/PKA signaling. We then performed a comprehensive pharmacology screen in hippocampal neurons and found that Gm38257, but not Arl5b, was upregulated by D1 receptor stimulation, KCl, and Glutamate stimulation, as well as to Forskolin treatment after TTX (Fig. S4B, Supplementary Table S1). Arc mRNA was used as a positive control for immediate early gene expression [1] and appeared to be expressed similarly to Gm38257 (Fig. S4B). Taken together, these results suggest that Gm38257 is regulated by activity, independently of Arl5b.

To confirm cAMP dependent dendritic transport and localization of Gm38257 in hippocampal neurons, we performed RNA FISH in stimulated and unstimulated conditions. By co-staining with Map2 to label dendrites and cFos to differentiate stimulated and unstimulated neurons, we confirmed that this lncRNA is globally upregulated and dendritically localized at 30 min and 60 min post stimulation [Somatic RNA: FSK 30 min vs DMSO 30 min: p=0.04, FSK 60 min vs DMSO 60 min: p = 0.03, Dendritic RNA: FSK 30 min vs DMSO 30 min: p = 0.02, FSK 60 min vs DMSO 60 min: p = 0.01, Pairwise two-tailed T-test with Holm correction, N = 17-24 neurons from 3 experiments] (Figs. 1F-H, Supplementary Table S2). We further observed that Gm38257 RNA puncta are distally localized in dendrites in a Forskolin-dependent manner (Fig. S5A). By quantifying the RNA puncta sizes into discrete bins (Fig. S5B,C, Supplementary Table S2), we learned that the increase in dendritic expression of Gm38257 was specific to large sized puncta (> 0.5 square microns), indicating a reliable signal difference between stimulated and unstimulated conditions. Furthermore, we fit a linear model to determine the relationship between nuclear cFos expression with somatic and dendritic Gm38257 expression. Indeed, cFos had a statistically significant positive correlation with both of these variables, indicating that cFos is a reliable predictor of Gm38257 expression (Fig. S5D). Consistent with our previous findings, Arl5b mRNA did not display activity-dependent dendritic localization (Figs. S6A-C). Malat1, an abundant nuclear enriched lncRNA, also showed no dendritic localization, as did the no probe control (Fig. S6D).

To confirm whether Gm38257 levels are transcriptionally regulated, we treated neurons with the transcription initiation complex inhibitor, ActinomycinD, prior to Forskolin treatment. Our FISH analysis shows that activity-dependent somatic and dendritic localization of Gm38257 is significantly reduced. (Fig. S7, Supplementary Table S3) in the presence of Actinomycin D, suggesting cAMP signaling induced transcriptional changes in Gm38257. These results are consistent with our previous data showing that Gm38257 is enriched in synaptoneurosomes as a nascent RNA (Fig. 1C). Conversely, blocking ribosome function with Anisomycin prior to Forskolin stimulation did not result in decreased somatic or dendritic expression of Gm38257 (Fig. S7, Supplementary Table S3). These results suggest that Gm38257 expression is dependent on transcriptional, rather than post transcriptional, processes.

RNA cargos that are synaptically targeted are known to require active transport along microtubule and actin cytoskeleton [22]. To confirm whether Gm38257 is likewise transported by cytoskeletal machinery, we blocked microtubule or actin cytoskeleton using inhibitors of polymerization Nocodazole or Cytochalasin D, respectively, prior to Forskolin treatment. We found that both of these manipulations reduced the dendritic, but not the somatic, expression of Gm38257, suggesting that this lncRNA is actively transported to dendrites and that integrity of microtubule and actin cytoskeleton is required for its localization (Fig. S8, Supplementary Table S3). This is consistent with previous findings regarding the transport of the small ncRNA BC1 [23]. Finally, by labeling dendritic spines with GFP and co-staining with Actin, the primary cytoskeletal component of dendritic spines, we observed colocalization of Gm38257 puncta with these spine markers, confirming that this lncRNA is localized to synaptic compartments (Fig. S9). Given that Gm38257 is acutely regulated by neuronal activity and is distally transported as a nascent RNA upon expression, we have termed this novel transcript ADEPTR—Activity DEPendent Transported lncRNA. Our results suggest that ADEPTR is an activity dependent intronic lncRNA that is expressed and distally transported to synapses independent of its host mRNA.

### lncRNA ADEPTR is required for activity-dependent changes in synaptic transmission and structural plasticity

We next investigated whether cAMP dependent changes in ADEPTR expression levels play a role in hippocampal neuron function by employing a loss of function analysis using antisense locked-nucleic acid Gapmers, a strategy that has previously been used to ascertain lncRNA function in neurons [11]. We transfected two independent Gapmers targeting ADEPTR, scrambled or no Gapmer for 48 hours into primary hippocampal neurons, then extracted RNA after treating neurons with Forskolin or DMSO (Fig. 2A). Both Gapmers significantly blunted the cAMP-dependent increase of ADEPTR in hippocampal neurons [Untransfected + FSK: 7.5±0.91, Scrambled Gapmer + FSK: 6.8±0.73, Gapmer1 + FSK: 3.7±0.67, Gapmer2 + FSK: 1.5±0.47, N=4-6 per group, p<0.05, One-Way ANOVA, Tukey’s HSD] (Fig. 2B, S10, Supplementary Table S4). Gapmer2 was also effective in suppressing ADEPTR levels in basal conditions [Untransfected + DMSO: 1.0±0.09, Scrambled Gapmer + DMSO: 0.98±0.13, Gapmer2 + DMSO: 0.24±0.05, p<0.05, One-Way ANOVA, Tukey’s HSD] (Fig. 2B, Supplementary Table S4). Activity-dependent expression of Arc mRNA was not affected by either Gapmer (Fig. S11A, Supplementary Table S4), suggesting no difference in cAMP-induced immediate early gene expression changes between samples. Although Arl5b mRNA was also downregulated by both Gapmers (Fig. S11B), Western blotting showed that at 30 minutes post stimulation, Arl5b protein levels were neither affected by ADEPTR Gapmers nor by Forskolin stimulation (Figs. S11C, D, Supplementary Table S4). We therefore conclude that the Gapmers are efficient in lowering cAMP-dependent ADEPTR expression without impacting Arl5b.

**Figure 2:**
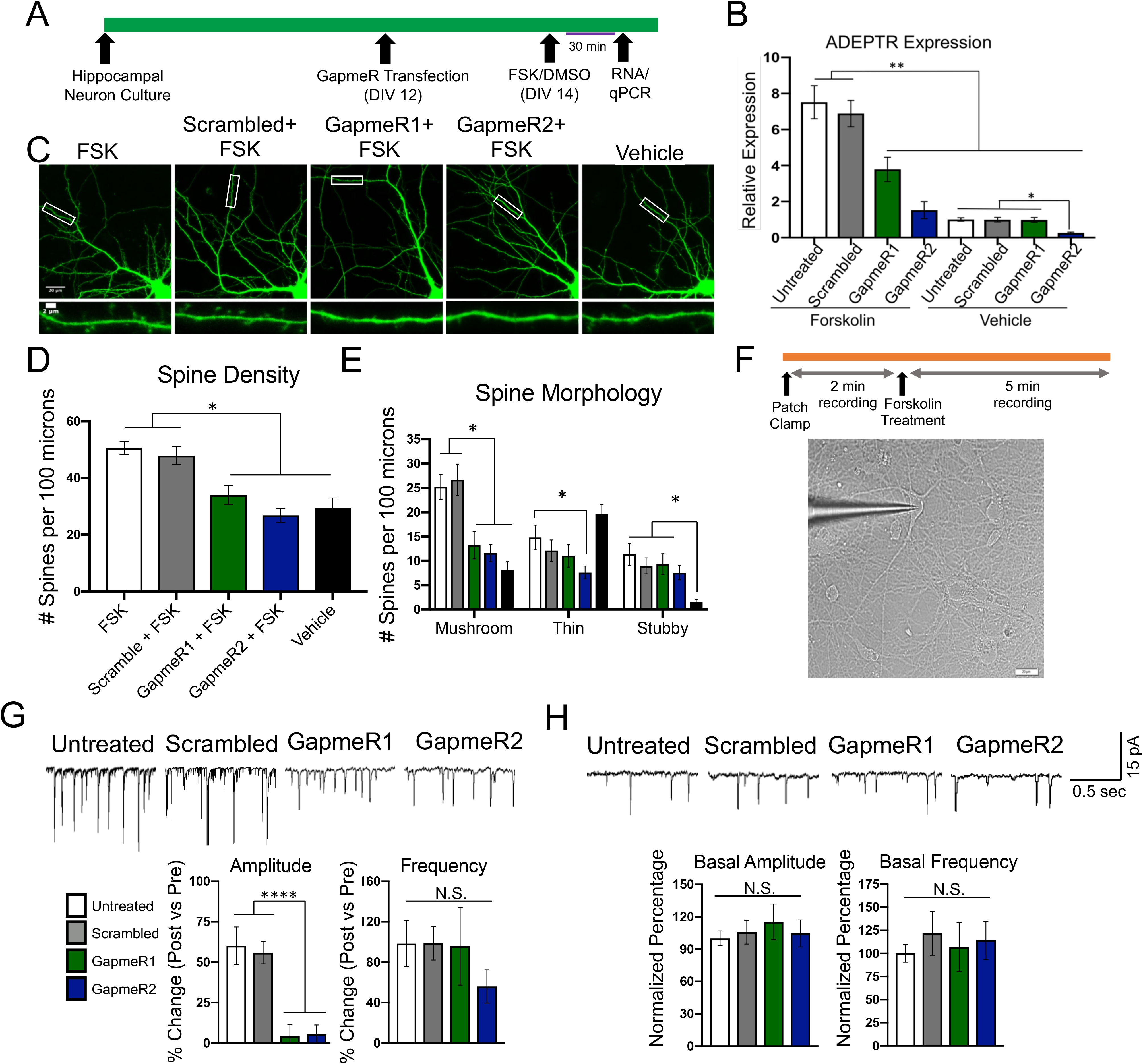
ADEPTR expression is required for cAMP-dependent changes in dendritic spine dynamics and synaptic transmission. **(A)** Experimental timeline of transfection for gene expression measurement. **(B)** qRT-PCR measuring ADEPTR expression in Forskolin or DMSO-treated hippocampal neurons after ADEPTR knockdown by two independent Gapmers. N=4-6 replicates per group. Error bars represent SEM. **p<0.01, *p<0.05. One-way ANOVA, Tukey’s HSD. **(C)** GFP labeled/Gapmer transfected cells were treated with Forskolin or DMSO for 30 min and subjected to live imaging. **(D)** Analysis of spine density and **(E)** morphology as performed by MATLAB custom script. (**F**) Top: Timeline of transfection for electrophysiology experiments. Bottom: Representative image of a patched neuron for sEPSCs recording in the voltage-clamp mode. Forskolin was added to the bath at a final concentration of 25uM. Scale bar is 20 microns. Analysis of amplitude and frequency from activity dependent **(G)** and basal **(H)** sEPSCs. N=22-28 neurons per group from 3 independent experiments. In all graphs, error bars represent SEM. **** p<0.0001, **p<0.01, One-way ANOVA, Tukey’s HSD.

Previous work has shown that cAMP signaling results in robust changes in synaptic transmission, as well structural plasticity, in hippocampal neurons [24]. We asked whether ADEPTR knockdown could influence neuron function in basal or cAMP-dependent conditions. We therefore performed live imaging of GFP-labeled hippocampal neurons in stimulated and unstimulated conditions and asked whether ADEPTR knockdown alters neuronal structural properties. By performing spine analysis, we discovered that both ADEPTR-targeting Gapmers suppressed cAMP-dependent changes in spine density [FSK+GFP: 50.59±2.3, FSK+ Scrambled Gapmer: 47.87±3.12, FSK+ Gapmer1: 33.95±3.32, FSK+ Gapmer2: 26.8±2.45, DMSO+GFP: 29.38±3.52, p<0.01, One-Way ANOVA, Tukey’s HSD, N= 16-22 neurons from 3 experiments] (Fig. 2C, D, Supplementary Table S5). Specifically, the deficiency in spine number was specific to mushroom spines [FSK+GFP: 25.22±2.5, FSK+ Scrambled Gapmer: 26.68±3.19, FSK+ Gapmer1: 13.24±2.84, FSK+ Gapmer2: 11.62±1.82, DMSO+GFP: 8.15±1.67, p<0.01, One-Way ANOVA, Tukey’s HSD] (Fig. 2E, Supplementary Table S5). Conversely, by performing Sholl analysis on these same neurons, we ascertained that neither ADEPTR knockdown nor cAMP signaling had an effect on neuronal branching (Fig. S12, Supplementary Table S5). We then measured the effect of ADEPTR knockdown on activity-dependent spontaneous excitatory post-synaptic currents (sEPSCs) using whole-cell patch-clamp recordings before and after Forskolin stimulation (Fig. 2F). Knockdown of ADEPTR by two independent Gapmers suppressed the cAMP dependent upregulation of EPSC amplitude [Untransfected: 60.16%±11.65, Scrambled Gapmer: 55.91%±6.95, Gapmer1: 4.12%±7.41, Gapmer2: 5.38±5.78, N = 22-28 neurons from 3 experiments] but not frequency [p=0.6, One-Way ANOVA] (Fig. 2G, Supplementary Table S6). Intriguingly, ADEPTR knockdown had no effect on basal EPSC amplitude or frequency suggesting its role is specific to cAMP dependent changes in synaptic transmission (Fig. 2H, Supplementary Table S6).

To confirm our findings by an alternative loss-of-function strategy, we cloned an shRNA cassette corresponding to the same sequence as Gapmer2 into a PLL3.7 shRNA plasmid with a lentiviral packaging backbone and GFP reporter. To confirm knockdown efficiency of this plasmid in neurons relative to scrambled shRNA plasmid, we packaged both into separate lentiviruses, transduced neurons, and collected RNA from FSK treated or untreated conditions. Like the Gapmer2, the ADEPTR shRNA significantly suppressed ADEPTR expression in basal and Forskolin conditions [shADEPTR vs shNC: p = 0.028, shADEPTR+FSK vs shNC + FSK: p = 0.017, Unpaired two-tailed T-Test, N=6-7 per group] (Figs. S13 A,B, Supplementary Table 7). However, unlike Gapmer2, Arl5b mRNA levels were unchanged [shADEPTR vs shNC: p = 0.31, shADEPTR+FSK vs shNC + FSK: p = 0.25, Unpaired two-tailed T-Test, N=6-7 per group] (Figs. S13 A,B, Supplementary Table 7). By imaging Forskolin or DMSO treated neurons expressing the shRNA or scrambled shRNA plasmid, and performing spine analysis, we confirmed that ADEPTR knockdown suppresses cAMP-dependent changes in total [shNC+FSK: 68.63±3.75, shNC+DMSO: 48.49±5.2, shADEPTR+FSK: 46.65±4.28, p <0.001, One-Way ANOVA, Tukey’s HSD] and mushroom spine density [shNC+FSK: 50.47±3.77, shNC+DMSO: 31.08±3.24, shADEPTR+FSK: 32.1±4.35, p <0.01, One-Way ANOVA, Tukey’s HSD, N = 19-31 neurons per group] (Figs. S13C-D, Supplementary Table 7). These data indicate that ADEPTR mediates activity-dependent changes in synaptic transmission and structural plasticity of dendritic spines.

### ADEPTR is required for transient structural plasticity in stimulated spines

Given that ADEPTR mediates activity dependent structural plasticity of dendritic spines and is transported to synapses, we hypothesized that ADEPTR specifically mediates the activity-dependent growth of stimulated spines. To test this hypothesis, we employed glutamate uncaging using two-photon excitation in organotypic hippocampal slice cultures [25] (Fig. 3A). We employed two independent shRNAs targeting ADEPTR, corresponding to Gapmer-1 and Gapmer-2 sequences, respectively, as well as a scrambled shRNA control. Neither shRNA targeting ADEPTR influenced immediate spine enlargement of uncaged spines at 0-70 seconds post uncaging (Figs. 3B,C, Supplementary Table 8). However, both ADEPTR-targeting shRNAs resulted in a transient reduction in dendritic spine size within 6.5 minutes of uncaging [1.5-3.5min: shNC: 150±15.5%, sh1: 107±14.72%, sh2: 70±8.97%; 4.5-6.5min: shNC: 104±18.7%, sh1: 55.1±12.1%, sh2: 51.94±8.3%, p<0.05, One-Way ANOVA, Dunnett’s Test, N =16-18 spines from 4-7 neurons per group] (Fig. 3D, E, Supplementary Table 8). Interestingly, there was no difference in sustained spine size by 20 minutes post uncaging between groups (Fig. 3E,F, Supplementary Table 8). Taken together, these results suggest that ADEPTR expression is critical to structural plasticity changes associated with stimulated dendritic spines.

**Figure 3:**
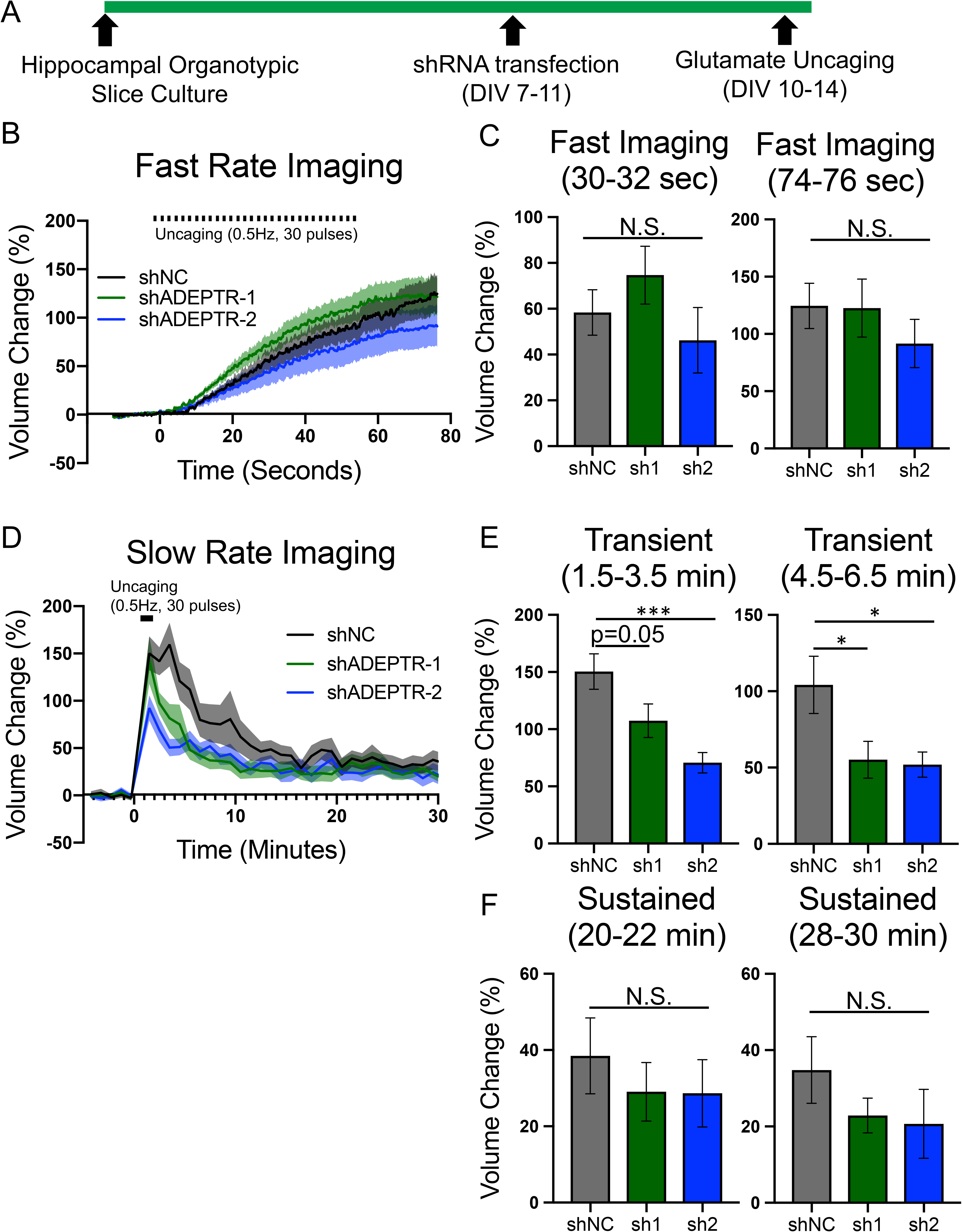
ADEPTR is required for transient structural plasticity in stimulated spines. **(A)** Experimental timeline of hippocampal organotypic slice culture and shRNA transfection for glutamate uncaging experiments. **(B)** and **(C)** analysis of rapid spine volume changes up to 1.5 min after uncaging. **(D)** Analysis of transient and sustained spine volume changes 1.5-30 min after uncaging. **(E)** Analysis of discrete time bins in the transient and **(F)** sustained phase of glutamate uncaging. *** p < 0.001, * p < 0.05. One-Way ANOVA followed by Dunnett’s Test, N =16-18 spines from 4-7 neurons per group.

### Activity-dependent ADEPTR transport is mediated by Kif2a

Given that ADEPTR is distally transported during cAMP signaling and mediates neuronal cAMP dependent changes in structural plasticity, we hypothesized that ADEPTR might carry out its function at or near the site of plasticity changes through binding partners. To test this hypothesis, we performed an RNA antisense purification (RAP) to pull down ADEPTR from Forskolin-treated hippocampal neurons (Fig. S14A) and analyzed protein and RNA interactors. This protocol involved the use of two long (∼300-400 nt) antisense probes on the 5’ and 3’ end of the putative ADEPTR transcript that sparsely contain biotin-incorporated Uracils for streptavidin-based pulldown (Fig. S14B). Sense probes were also implemented to control for non-specific binding. qRT-PCR analysis showed that the RAP protocol robustly and specifically enriched ADEPTR [Fold change 21.75±3.07 relative to sense control, N=5-8 per group, p < 0.05, Unpaired Two-tailed T-Test]. Notably, neither Arl5b nor Actin mRNA were preferentially enriched, indicating that the RAP protocol is specific to ADEPTR enrichment (Fig S14C).

We first examined protein component of ADEPTR pull-down complexes and searched for molecular motors that might mediate ADEPTR localization. Using LC-MS/MS analysis, we identified a unique protein enriched in the ADEPTR RAP complex: Kinesin motor Kif2a (Fig 4A). We validated this interaction through repeated independent pulldown experiments, via Western blotting [N=4-7 per group from 3 experiments, p<0.05, One-Way ANOVA, Tukey’s HSD] (Fig 4B). As an additional control we also incorporated proteins enriched in Malat1 RAP complexes. Given that Kif2a is part of the Kinesin family of motor proteins responsible for anterograde transport, we hypothesized that Kif2a mediates ADEPTR transport. To test this hypothesis, we knocked down Kif2a, or 4 other Kinesins, using siRNAs that were previously validated in hippocampal neurons for knockdown efficiency [26]. Following 72 hours of knockdown, we treated neurons with Forskolin prior to synaptoneurosome fractionation or ADEPTR FISH (Fig. 4C). qRT-PCR analysis from synaptosomes showed that Kif2a siRNA significantly downregulated ADEPTR expression in synaptoneurosomes relative to scrambled siRNA [ADEPTR Fold change in siKif2a relative to siNC: 0.45±0.18, N=3-4 per group. p<0.05, Dunnett’s Test]. Importantly, knockdown of Kif17, Kif21a, Kif11, and Kif5c did not show a concomitant decrease in ADEPTR expression. Additionally, Arl5b and Arc mRNA expression in synaptosomes were not affected by any siRNA tested (Fig. 4D, Supplementary Table 9). We then repeated Kif2a knockdown and measured ADEPTR expression using FISH (Fig. 4E). While the somatic fluorescence intensity of the ADEPTR channel was not significantly impacted (Fig. 4F), the dendritic puncta account was significantly reduced, though not entirely suppressed, in Kif2a deficient neurons [siNC: 127.28±15.8, siKif2a: 81.14±7.52, N=25-28 neurons per group from 3 independent experiments, p<0.05, Welch t-test] (Fig. 4G, Supplementary Table 9). Taken together, these results show that activity-dependent dendritic localization of ADEPTR is mediated by Kif2a.

**Figure 4:**
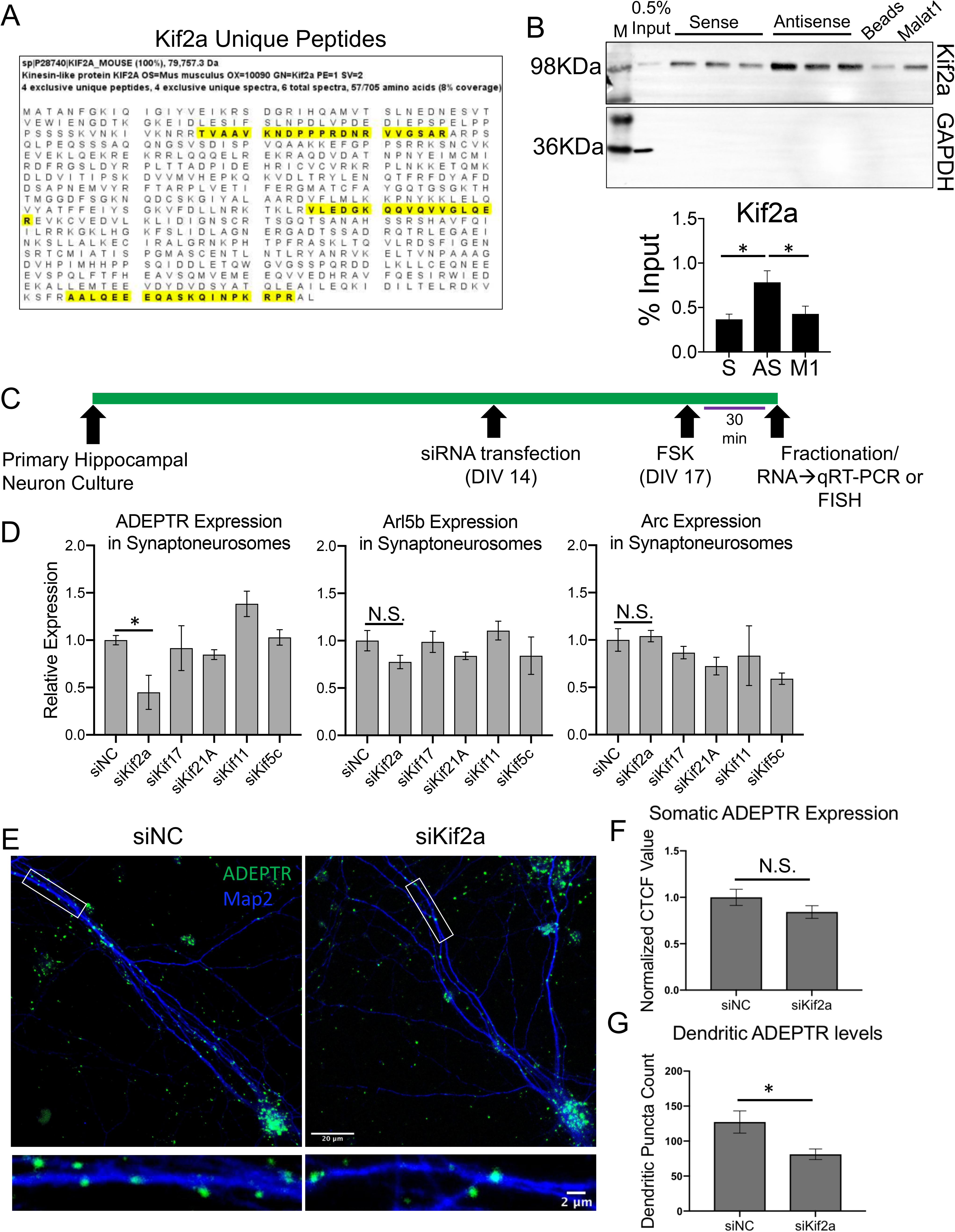
Activity-dependent ADEPTR transport is mediated by Kif2a. **(A)** (Top) Unique peptide counts of Kif2a from ADEPTR Antisense Purification, identified by LC-MS/MS. (Bottom) Sequence coverage, percent chance of correct protein identification, and number of unique peptides and spectra. **(B)** (Top) Western blotting of ADEPTR pulldown. Sense probes (S), beads alone, and Malat1 (M1) were used as controls. (Bottom) Quantification of Western blotting based on % input. N=4-7 per group from 3 experiments. * p<0.05. One-way ANOVA, Tukey’s HSD. **(C)** Timeline for RNAi-mediated knockdown of candidate Kinesins for fractionation or FISH studies. **(D)** Neurons were treated with Forskolin prior to fractionation and synaptosome RNA extraction. Graph shows qRT-PCR analysis of ADEPTR, Arl5b, and Arc expression in Kif2a, Kif17, Kif21A, Kif11, and Kif5c siRNA groups, relative to control. N=3-4 per group. *p<0.05, Dunnett’s Test. **(E)** FISH images from siKif2a+FSK or siNC+FSK. **(F)** Quantification of somatic and **(G)** dendritic ADEPTR expression. N=25-28 neurons per group from 3 independent experiments. **p<0.001 Student’s T-test. Error bars represent SEM. Scale bar represents 20 microns in the neuron image and 2 microns in the inset.

### ADEPTR interacts with the Spectrin/Ankyrin complex and regulates distal AnkB localization

LC-MS/MS analysis of ADEPTR RAP identified two additional associated proteins: Sptn1 and AnkB. We validated this interaction through independent pulldown experiments, via Western blotting [AnkB and Sptn1: p<0.05, One-Way ANOVA, Tukey’s HSD, N=4-7 replicates from 3 experiments] (Fig. 5A, B, Supplementary Table 10). It is known that early to mid-stage plasticity changes are mediated by post translational activities such as protein-dependent signaling cascades resulting in cytoskeletal rearrangements [27]. Sptn1 (Alpha-II-Spectrin) is a filamentous protein that interacts with its complementary beta subunit to scaffold the actin cytoskeleton [28]. It exhibits a unique pattern of periodicity in neuronal processes that complements periodic actin [29, 30]. Importantly, Spectrin is anchored to the plasma membrane through Ankyrin proteins. AnkB (Ank2) is a brain-enriched Ankyrin member that is expressed in both axons and dendrites [31]. Together, the Spectrin/Ankyrin complex has been known to be required for Actin-enriched spine formation [32] and neurite assembly [33]. Sptn1 and AnkB were also shown to interact in neurons to stabilize the T-type calcium channel Cav3.2 [34]. However, it remains unresolved how this network is regulated in neurons.

**Figure 5:**
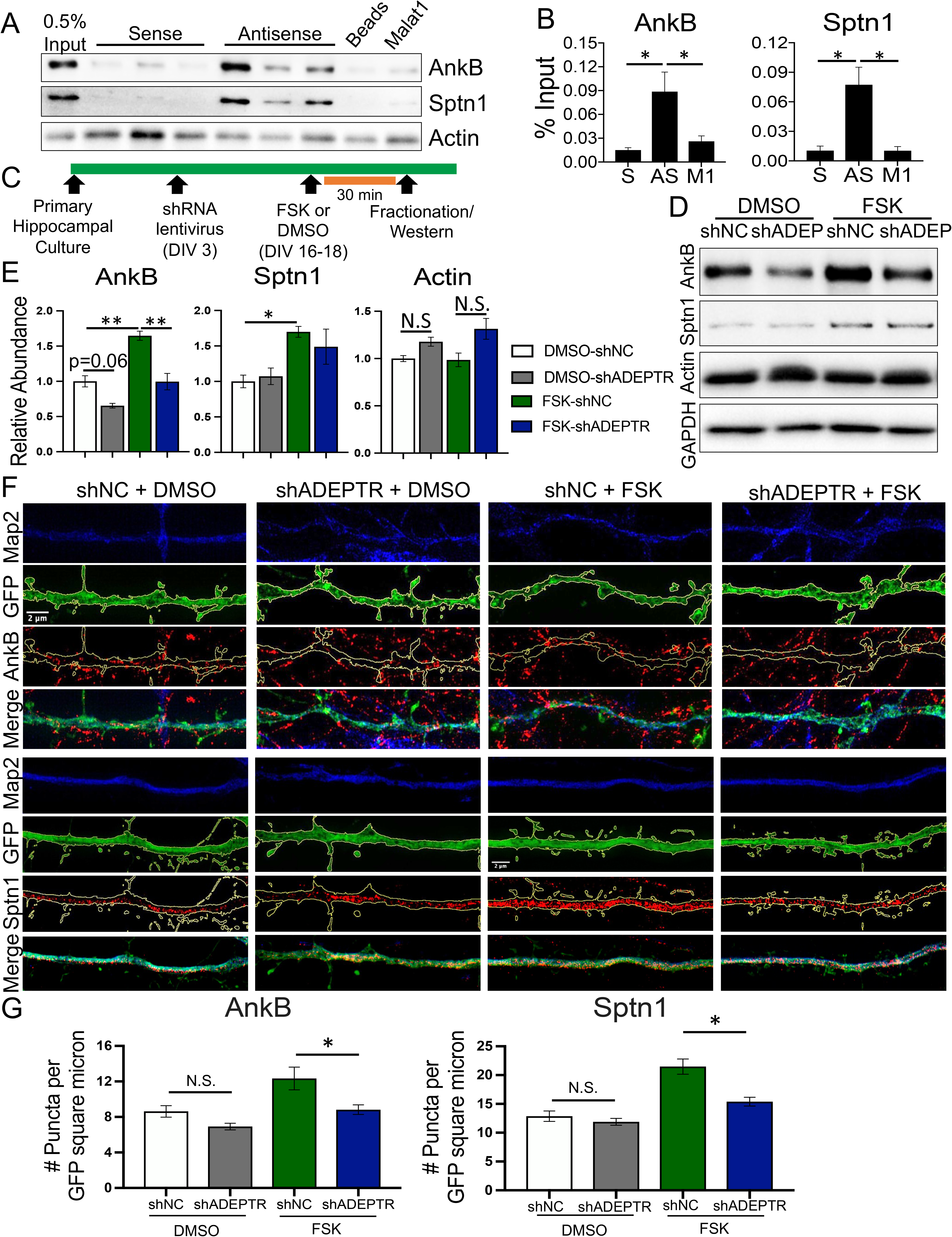
ADEPTR binds to Sptn1 and AnkB and regulates their activity-dependent dendritic localization in hippocampal neurons. **(A)** Western blotting of ADEPTR Antisense Purification. Sense probes (S), beads alone, and Malat1 (M1) were used as controls. **(B)** Quantification of Western blotting based on % input. N=4-7 per group from 3 experiments. * p<0.05, One-way ANOVA, Tukey’s HSD. **(C)** Timeline for RNAi-mediated lentiviral knockdown of ADEPTR for synaptoneurosome fractionation studies. **(D)** Western Blotting of indicated proteins in synaptoneurosome fractions. **(E)** Quantification of AnkB, Sptn1 and Actin expression in synaptoneurosomes N=4-6 per group. *** p<0.001, *p<0.05, Two-way ANOVA, Tukey’s HSD. **(F)** (Top) SIM images of AnkB and (Bottom) Sptn1 within GFP-labeled neuronal processes from shNC or shADEPTR expressing FSK-treated neurons. **(G)** Quantification of respective protein puncta. N=30-38 dendrite sections from 16-22 neurons per group. *p<0.05 Two-Way ANOVA, Tukey’s HSD. Error bars represent SEM. Scale bar represents 2 microns.

Given that ADEPTR regulates activity-dependent spine changes in mature hippocampal neurons, we asked whether ADEPTR might regulate components of the Spectrin/Ankyrin network. Therefore, we globally knocked down ADEPTR in hippocampal neurons using a lentivirus expressing ADEPTR shRNA, or control, and isolated protein from synaptoneurosomes after treating neurons with Forskolin or DMSO (Fig. 5C). We found, by Western Blotting, that both Sptn1 and AnkB protein expression in synaptoneurosomes is upregulated by cAMP signaling. Interestingly, AnkB, but not Sptn1, expression was reduced in ADEPTR-deficient neurons, in a cAMP-dependent manner in synaptoneurosomes [AnkB: shNC+FSK vs shADEPTR+FSK: p<0.01, Two-Way ANOVA, Tukey’s HSD, N=4-6 per group] (Fig. 5D, E). Further, Actin protein levels were not affected by either FSK treatment or ADEPTR knockdown (Fig. 5D, E).

To better understand the role of ADEPTR in AnkB and Sptn1 expression in neuronal distal processes, we sparsely labeled neurons with the shRNA plasmid, and performed super resolution imaging of GFP labeled distal neuronal processes immuno-stained with AnkB or Sptn1 under cAMP-conditions (Fig. 5F). Structured Illumination Microscopy of distal neuronal processes confirmed that ADEPTR knockdown suppresses the dendritic localization of AnkB and Sptn1 only under cAMP conditions [AnkB and Sptn1: shNC+FSK vs shADEPTR+FSK p<0.05, Two-Way ANOVA, Tukey’s HSD, N=30-38 dendrite sections from 16-22 neurons from 3 experiments] (Figs. 5G, H). These results indicate that ADEPTR mediates dendritic localization of AnkB and Sptn1 thereby facilitating structural changes at the synapse.

### A 222-nucleotide region of ADEPTR interacts with AnkB and Sptn1

We then asked what regions of ADEPTR are directly involved with protein binding. To address this question, we employed a modified version of the RNA Antisense Purification strategy that draws from elements of ribosome foot-printing [35] that has previously been employed to study lncRNA interactions [36]. Briefly, Forskolin-treated mature hippocampal neurons were chemically crosslinked, lysed, and sonicated. Upon pulldown of ADEPTR, unprotected regions of ADEPTR in the RAP complex were broken down by RNAse treatment and separated from the protected fragments bound by protein using sucrose-cushioned ultracentrifugation. The RNAse-protected RNA in the precipitate was then isolated for further analysis (Fig. 6A). Compared to the sense probe control, the RNA profile of the RNAse-protected ADEPTR RAP complex showed a more robust distribution of peaks, which concentrated in the 100-400nt range (Fig. S14D). We then performed RNAseq and mapped the reads to the ADEPTR locus. Surprisingly, we identified one predominant fragment at the 3’ end of the putative transcript that had a length of 222nt (Fig. 6B). We then performed a pulldown from hippocampal neuron lysates using biotinylated oligo probes of an extended version of this sequence (position 2415-2700 of the putative ADEPTR) or previously used sense probe (1924-2221) (Fig. 6C). Beads alone were used as a control for background levels of enrichment. We found that this sequence is sufficient to enrich AnkB and Sptn1, but not Actin or GAPDH, suggesting that this sequence is sufficient to pulldown the AnkB and Sptn1 [AnkB: 2415-2700 vs Beads: p<0.01, Sptn1: 2415-2700 vs Beads: p<0.05, One-Way ANOVA, Dunnett’s Test compared to Beads, N=3-5 replicates from 3 experiments] (Fig. 6D,E, Table S11).

**Figure 6:**
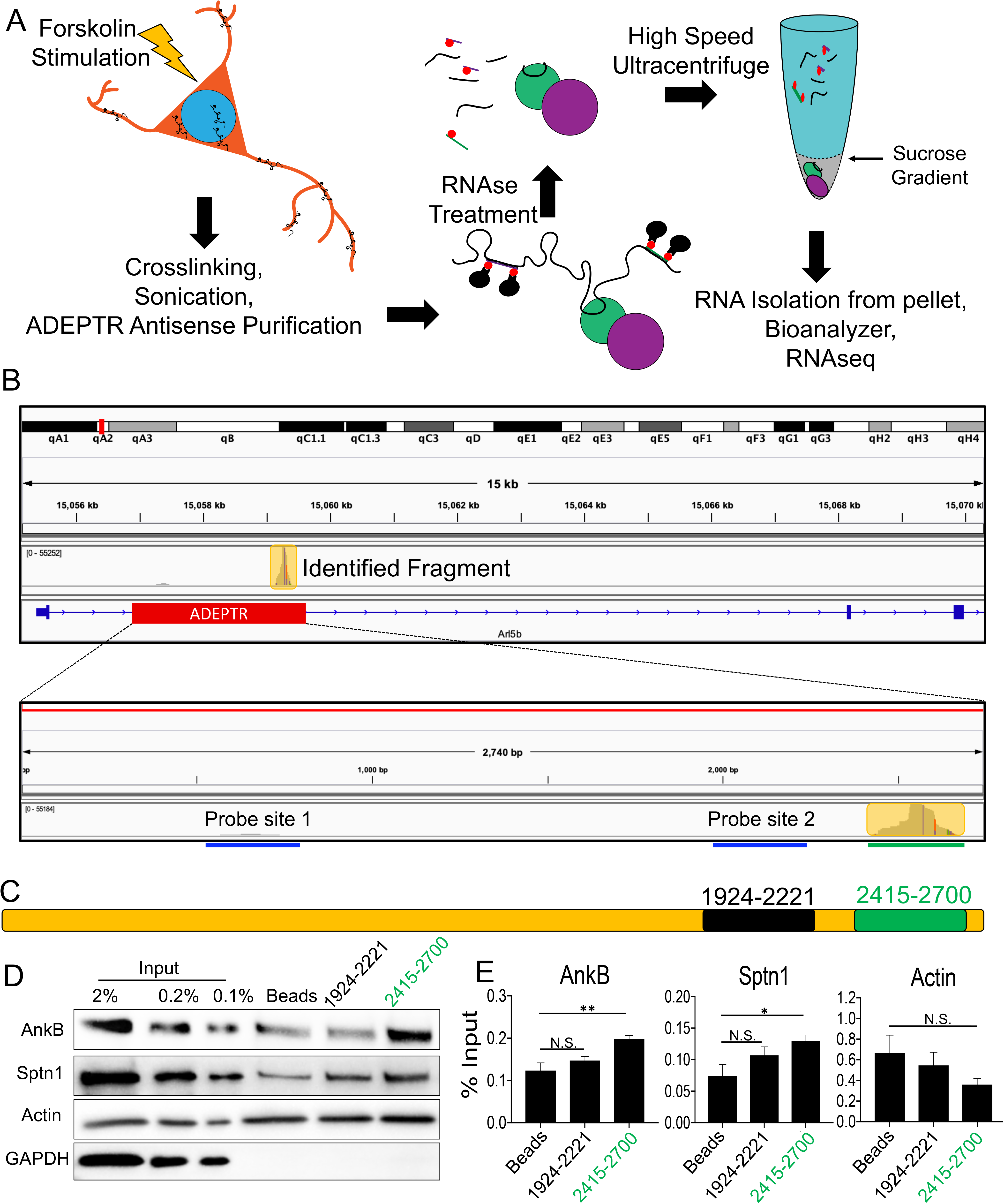
Identification of protein binding sequences of ADEPTR. **(A)** Schematic summarizing the strategy to pulldown ADEPTR under crosslinked conditions, purify the protected regions using RNAse treatment and ultracentrifugation under sucrose gradient, and identification of fragments by RNAseq. **(B)** Reads were mapped to the mouse genome using STAR and transcriptome using Salmon and visualized on IGV software. **(C)** Schematic of regions of ADEPTR used for making sense probes. Green probe represents the conserved domain and black represents the same sense probe control used in Figures 4 and 5. **(D)** Western blotting and **(E)** quantification of RNA pulldowns using the ADEPTR fragments. N=3-5 per group, from 3 experiments. **p<0.01, *p<0.05, One-Way ANOVA, Tukey’s HSD. Error bars represent SEM.

### ADEPTR directly interacts with the Ankyrin/Spectrin network via a 32-nucleotide sequence found in several human mRNA 3’UTRs

Cross-referencing this fragment with known features of the mouse reference genome, we discovered that it contains multiple conserved regions, as well as a short-interspersed nuclear element (SINE) (Fig. 7A). We then performed BLAST analysis on this fragment to the human genome and noted a ∼32nt region that mapped to multiple human mRNAs—most strongly to the negative transcription elongation factor NELFCD, peroxisome biogenesis factor PEX26, cation channel subunit TRPC3, and the glycosylphosphatidylinositol anchor protein PIGT. Notably, for all mRNAs analyzed by E-value thresholding, this region was shown to reside within their 3’UTRs (Fig. 7B). We hypothesized that this fragment within ADEPTR is specifically responsible for directly binding of Sptn1, AnkB, or both. Therefore, we prepared three separate constructs for generating biotinylated probes: a 837nt sequence corresponding to the 3’ end of the putative ADEPTR sequence (WT), the same construct lacking the 32nt binding region (Del), and one containing the reverse complement of the binding region (Sub) (Fig. 7C). Western blotting of pulldowns from hippocampal neuron extracts showed that the WT fragment had a significantly higher enrichment of both AnkB and Sptn1 compared to Beads alone, while neither the Del nor Sub fragments had a significant enrichment of these proteins [AnkB and Sptn1: WT vs Beads: p<0.001, One-Way ANOVA, Dunnett’s Test, N=3 replicates from 2 experiments] (Fig. 7D,E, Table S11). Actin and GAPDH were not preferentially enriched in any pulldown condition.

**Figure 7:**
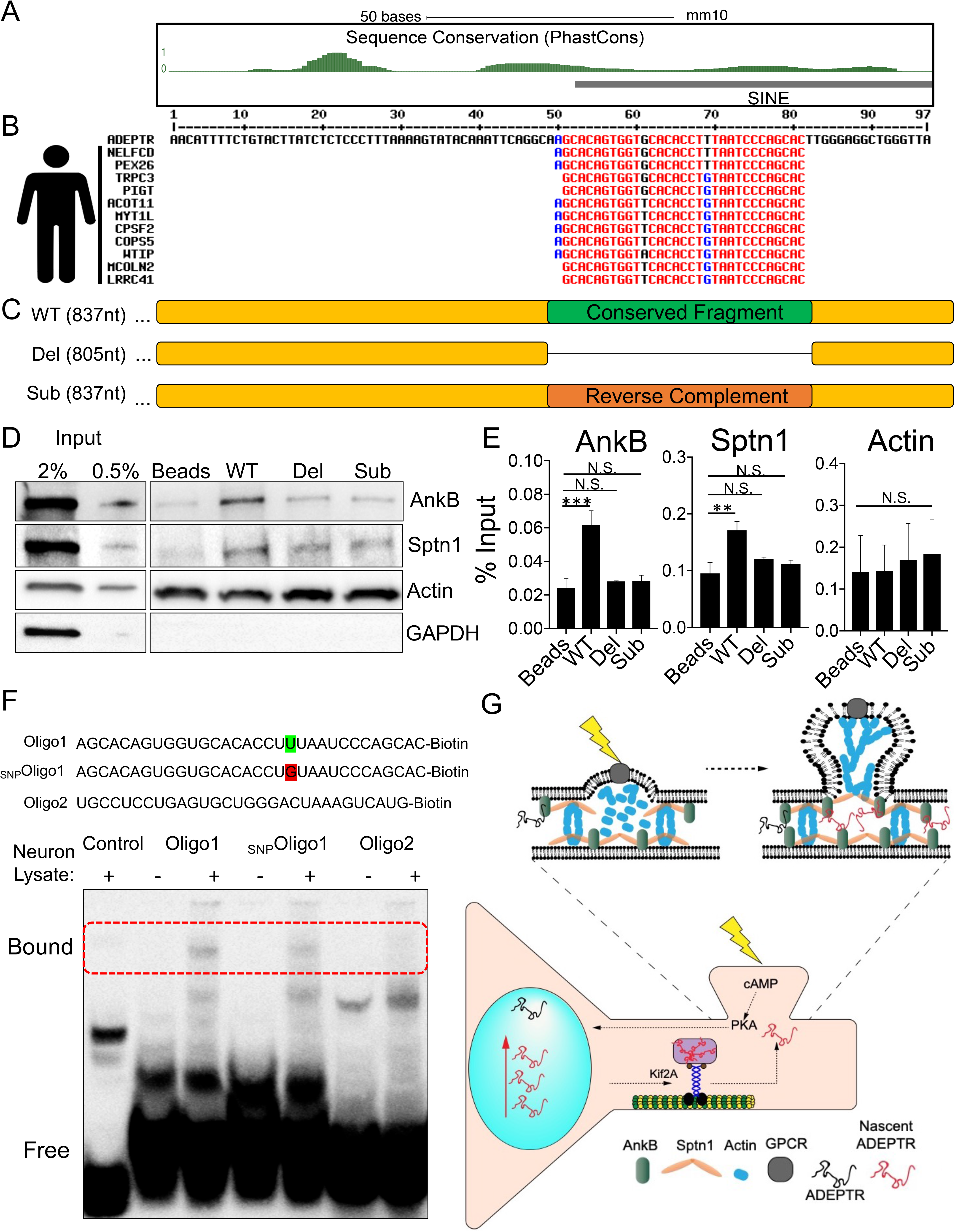
ADEPTR directly interacts with the Ankyrin/Spectrin network via a conserved domain found in human mRNA 3’UTRs. **(A)** The fragment identified was mapped to UCSC genome browser and analyzed for conservation (PhastCons) and presence of repeat elements (RepeatMasker). **(B)** A 97nt region showing highest conservation was Blasted to the human cDNA reference using Ensembl, and top hits (E-value < 0.001) were aligned using Multalin. Mapping to each human mRNA 3’UTR was performed using Ensembl Biomart. **(C)** Schematic of biotin-labeled fragments of ADEPTR containing the conserved region, a deletion, or substitution. **(D)** Western blotting and **(E)** quantification of RNA pulldowns using the ADEPTR fragments shown in **(C)**. N=3 per group, from 2 experiments. **p<0.01, *p<0.05, One-Way ANOVA, Tukey’s HSD. Error bars represent SEM. **(F)** (Top) Sequences of short oligos corresponding to the 32nt conserved domain (Oligo1), a single point mutation (_SNP_Oligo1) or another region of ADEPTR (Oligo2). (Bottom) RNA electrophoretic mobility shift assay of the three oligos, plus IRE control, with or without the presence of neuronal lysate. **(G)** Proposed model for ADEPTR in mediating structural plasticity.

Lastly, to confirm the significance of this conserved region in binding to protein via an alternative strategy, we asked whether the 32-nucleotide fragment alone could interact with proteins. Therefore we performed an RNA electrophoretic mobility shift assay (EMSA) on hippocampal neuron extracts using 3’end biotinylated conserved 32nt RNA sequence (Oligo1) of ADEPTR, the same construct with a single point mutation (_SNP_Oligo1), an alternative region of ADEPTR (Oligo2), or IRE Control RNA. We observed that protein extracts from hippocampal neurons bind Oligo1 better than the other 3 RNAs tested (Fig. 7F) suggesting functional interactions of the conserved 32 nucleotide region. Taken together, these data show that ADEPTR directly interacts with the Ankyrin/Spectrin network via a 32 nucleotide repeat element that is also found in 3’UTRs of few human mRNAs.

## Discussion

Synaptically localized RNAs are an important component of neuronal function and cognitive processes such as long term memory [37] but much less is known about the identity of noncoding components, regulation of their localization and function. Previous works have attempted to unearth the synaptic transcriptome through multiple approaches, such as Kinesin immunoprecipitation [38], neurite sectioning [39, 40], and synaptosome fractionation [41], coupled with micro-array or RNA sequencing. Our analyses, combining next generation RNA sequencing, qRT-PCR, FISH, and nascent RNA capture have identified a novel noncoding RNA transcript—ADEPTR—as an intronic lncRNA that is synaptically targeted in a cAMP-dependent manner, independent of its protein-coding host gene. It has been previously reported that functional intronic sequences can be employed as retained introns within mRNAs [42, 43], excised products from pre-mRNAs [44, 45], and as stable sequences with independent transcription [46]. Many of these sequences can be found in the cytoplasm [45]. In line with the latter, our work has shown that ADEPTR is an independently expressed and transported RNA, relative to its protein-coding host gene. Through pharmacology screening, we identified that ADEPTR is specifically regulated by excitatory signaling, and unlike Arl5b, its cAMP-induced expression is dependent on the activity of PKA.

Loss of function experiments, utilizing LNA Gapmers and shRNAs— have established that this lncRNA is necessary for cAMP-dependent structural plasticity of dendritic spines in hippocampal neurons, but not overall dendritic morphology. Supporting electrophysiology experiments showed that ADEPTR does not influence synaptic transmission under basal conditions, but rather specifically mediates cAMP-dependent changes in EPSC amplitude. Finally, glutamate uncaging experiments showed that ADEPTR is required for activity-dependent transient spine enlargement specifically at stimulated spines. Though previously identified lncRNAs have shown an influence in basal hippocampal neuron function—these include Malat1, NeuroLnc, NEAT1, and Gm12371 [11, 47–49]—none of these lncRNAs are targeted to synapses in an activity dependent manner for regulating spine dynamics. Given the discrete expression pattern of ADEPTR under cAMP signaling, our results suggest that ADEPTR plays a defined, yet potent role in activity-dependent structural plasticity (see Fig. 7G).

We next found that the distal localization, but not the expression, of ADEPTR is dependent on microtubule and actin polymerization. This suggested that ADEPTR is being actively trafficked by canonical transport mechanisms, of which Kinesins play a major role. In line with this observation, we have uncovered a specific Kinesin motor— Kif2a—as a binding partner and mediator of ADEPTR distal transport. Kif2a is an M-terminal Kinesin and, since its discovery, has been shown to carry cargoes in neurons [50, 51]. It has also been shown to function as a microtubule destabilizer, regulating neuronal outgrowth [52]. However, its microtubule destabilizing function can be inhibited by BDNF signaling, suggesting the possibility of a dual role for this protein that is regulated by activity [52]. Future study will require a deeper molecular understanding of how different converging signaling pathways result in Kif2a-dependent distal localization of ADEPTR, and other potential cargoes.

We then showed that ADEPTR binds to the Spectrin/Ankyrin network, specifically Sptn1 and AnkB. Furthermore, the interaction of ADEPTR with this network is functional as its expression is necessary for the cAMP-dependent localization of AnkB in synaptoneurosomes and both AnkB and Sptn1 in dendritic processes. Importantly a 222nt fragment of ADEPTR is sufficient to enrich AnkB and Sptn1. Previously discovered lncRNAs have been shown to interact with proteins with no defined RNA-binding capabilities, through protected domains [46, 53], demonstrating an intriguing capability of regulatory RNAs typically ascribed to proteins. Here, we found that the interaction within this protected region is through a 32 nucleotide SINE B1 domain, found in 3’ UTRs of human mRNAs, which we confirmed through deletion and substitution experiments. It’s previously been shown that a lncRNA functions in the cytoplasm of dopaminergic neurons through a SINE ALU repeat to salvage the expression of an mRNA [36], though the functional consequence of this rescue in neurons warrants further study. Furthermore, a brain-enriched zebra fish lncRNA lacking direct homologs in higher organisms has previously been shown to regulate zebra fish behavior via interaction with miR-29. Importantly, its miRNA interaction site is conserved in the 3’UTR of an mRNA in mice and humans [54]. Interestingly, computational analysis has drawn parallels between lncRNAs and 3’UTRs in terms of structural features and sequence composition, namely that they have similarly low protein coding potentials, have a low GC content, and show a less stable secondary structure compared to that of ORFs and even 5’UTRs. These suggest that both lncRNAs and 3’ UTRs are more reliant on protein interactions for their stability than mRNAs [55]. In line with these discoveries, our findings show that ADEPTR functions in the cytoplasm through cytoskeletal-regulating proteins AnkB and Sptn1, and is subsequently turned over. The identification of a conserved functional region of ADEPTR that matches to multiple human mRNA 3’UTRs is further evidence that noncoding elements—even within protein-coding genes—can play a role in structural plasticity. Indeed, ADEPTR activity seems to fit this pattern of regulation and may put fourth an intriguing role for lncRNAs locally regulating sites of structural plasticity.

Although Ankyrin is a canonical binding partner of Spectrin subunits, the precise mechanism of how these proteins regulate structural plasticity remains to be elucidated. Indeed, AnkB is known to bind a multitude of distal targets including the plasma membrane, different Spectrin isoforms, L1-CAMs and Calcium channels [37, 38]. Given the previously established role for AnkB in regulating voltage-gated Calcium channels in dendrites [34], it is possible that by scaffolding AnkB, ADEPTR is indirectly involved in mediating Calcium influx for activity-dependent remodeling of synapses.

lncRNAs are emerging as subtle, yet powerful regulators of gene expression and cellular function, and their subcellular localization is an important indicator of their function. Akin to previously studied lncRNAs [56] ADEPTR shows both a nuclear and somatic expression upon activity, thereby raising an important question as to how it carries out its phenotype. Given the rapid expression and shuttling of ADEPTR to distal processes, as well as the interaction with structural proteins Sptn1 and AnkB, our results suggest that ADEPTR carries out its function predominantly in the distal cytoplasm. Cytoplasmic lncRNAs have also been demonstrated to act as scaffolds for protein complexes: a role also attributed to UTRs or non-coding isoforms of mRNAs [57–59].

Our study is the first to describe a cAMP/PKA-dependent lncRNA that is expressed, distally transported by a kinesin motor protein, and regulates structural plasticity through functional interactions with the Spectrin/Ankyrin network. This study illuminates our understanding of non-coding elements of the synaptically targeted transcriptome, of which we know very little, and puts forth synaptically localized lncRNAs as important regulators of adaptive neuronal function.

## Materials and Methods

### Animals

CD1 pregnant mice (Charles River Laboratories) were housed individually on a light dark cycle (12 hour/ 12 hour) with ad libitum access to food and water. Experiments were performed during the light part of the diurnal cycle. Housing, animal care and experimental procedures were consistent with the Guide for the Care and Use of Laboratory Animals and approved by the Institutional Animal Care and Use Committee of The Scripps Research Institute.

#### Primary Hippocampal Neuron Culture

Hippocampi were dissected from E18 CD1 pups and plated on poly-D-lysine coated plates in Neurobasal media supplemented with Glutamax, Penstrep, and 5% FBS. Density of neurons plated was determined by each experimental requirement. Four hours after plating, media was replaced with feeding media consisting of Neurobasal media supplemented with Glutamax, Penstrep, and 2% B27 (Invitrogen), and half of the media was replaced every 4 days until the time of experiments at 37°C with 5% CO_2_.

### Transfection of plasmid constructs, LNA Gapmers, and siRNAs

GFP reporter plasmids (eGFP, PLL3.7 shRNA) were transfected into neurons at DIV 14 using Lipofectamine 2000 (Invitrogen). LNA Gapmers targeting ADEPTR or siRNAs targeting kinesins were transfected using Lipofectamine RNAiMax (Invitrogen). LNA Gapmers were synthesized by Exiqon (now QIAGEN) and transfected as previously described [15]. Briefly, siRNAs and Gapmers or shRNAs were introduced to primary hippocampal neurons (5–8 days *in vitro* (DIV)) using Lipofectamine RNAiMAX or Lipofectamine 2000 (Invitrogen) according to manufacturer’s guidelines. One day before transfection, the fresh culture medium was prepared and mixed evenly with the old medium. One-half of the mixed media were added to the cells for transfection, and the other one-half was saved for medium replacement for transfection. The sequence for Gapmer1 targeting ADEPTR is: 5’ TTAGGTGCCAGAATAA 3’. Gapmer2 targeting ADEPTR: 5’ TTTCCATCTTGCACCTAGCAG 3’. Scrambled Gapmer: 5’ AACACGTCTATACGC 3’. siRNAs targeting kinesins were previously used and validated [30]. The sequences for shRNA 1 and 2 cassettes for ADEPTR knockdown correspond to the Gapmer 1 and 2 sequences, respectively. Scrambled shRNA cassette: 5’AGTTCCAGTACGGCTCCAA3’.

### Pharmacology

Additional pharmacological agents used on hippocampal neurons were PKA inhibitor 14-22 amide (2.5uM), A2A receptor agonist Adenophostin A (5nM), PKC activator PMA (Phorbol 12-myristate 13-acetate, 1uM), D1 Receptor Agonist SKF38393 (10uM), L-Glutamate (500 nM), Potassium Chloride (50mM), and tetrodotoxin (TTX, 1uM).

### Cell Fractionation

For Synaptoneurosome preparation, DIV 16-18 hippocampal neurons were washed once with ice cold PBS and lysed in Syn-PER Synaptic Protein Extraction Reagent (Thermofisher), as per manufacturer’s instructions. For nuclear/cytoplasmic isolation, neurons were washed with cold PBS and lysed in cytoplasmic buffer (Tris-HCl 20mM, KCl 150mM, MgCl_2_ 1.5mM, DTT 1mM, 0.5% NP-40, and 1× EDTA-free protease and phosphatase inhibitor cocktails) and gently centrifuged at 4°C to separate the nuclear fraction (pellet) from the cytoplasmic supernatant. Both fractions were mixed with Trizol for RNA isolation and qRT-PCR.

### Quantitative Real-Time PCR

RNA was isolated using Trizol and reversed transcribed into cDNA using qScript cDNA SuperMix (Quanta Bio). qRT-PCR was performed in 384 well plates using SYBR Green master mix (Applied Biosystems) for detection in the ABI 7900 thermal cycler (Applied Biosystems). dCT values were obtained using Ct values for 18s for normalization, followed by relative quantification using 2^-ΔCT^ method.

### RNA Sequencing and Analysis

Total RNA sequencing was performed following the Scripps Florida Genomics core protocol, as described previously [11]. Fastq files were first assessed for quality using fastqc and trimmed using Trim Galore on the Scripps high-performance computer cluster. Fastq reads were then aligned to the *mus musculus* transcriptome using Salmon [60], also using Scripps high-performance computing cluster, to generate raw counts and TPM data based on Ensembl Transcript IDs. Data was then imported into R using tximport Bioconductor package and clustered using clusterProfiler. Differential expression analysis was performed using DEseq2.

### Nascent RNA capture

Nascent RNA capture was performed using the Click-iT Nascent RNA Capture Kit (Life Technologies) using manufacturer instructions. Primary hippocampal neurons were treated with 5-ethynyl Uridine at DIV 16. 24 hours later, neurons were treated with Forskolin (50uM) or DMSO (equal volume) for 30 minutes then lysed in Trizol for total RNA extraction. 10% of the total RNA was saved while the remaining 90% was processed for click-chemistry based azide-biotinylation of the 5-EU, followed by binding to MyOne Streptavidin T1 beads and elution.

### RNA FISH and immunofluorescence

Two 300-400 bp fragments of ADEPTR, Arl5b, and Malat1 cDNA were sub cloned into PCRII TOPO vectors and in vitro transcribed into DIG labeled probes for fluorescence in situ hybridization (FISH). Anti-DIG fab fragment antibody (Roche) was used at a concentration of 1:4000, and Tyramide signal amplification (ThermoFisher) was used using manufacturer instructions. One of the fragments, targeting the 3’ end of ADEPTR, was more efficient at producing a robust signal, and therefore was used for all subsequent studies. Immunofluorescence was performed in conjunction to or separate from FISH, according to experiment and samples were imaged on the Zeiss LSM 880 confocal microscope at Max Planck Florida Institute light microscopy facility. Neurons were plated at a low density (80,000 cells per well of a 24-well plate) to allow for imaging of single neurons at a time, with minimal crossover from other neurons.

### FISH puncta acquisition and analysis

To analyze dendritic RNA puncta count, for each neuron imaged, the Map2 labeled dendrites were traced in Fiji ImageJ (NIH) and FISH puncta were acquired using a custom ImageJ macros script. The puncta count per neuron and the size of each punctate (in microns) were exported into R for statistical analysis. The puncta sizes were divided into discrete bins in R and the frequency per bin was quantified for each experiment.

### Spine and Sholl analysis

Spine and Sholl analysis were performed as previously described [11]. Briefly, GFP expressing neurons were brought to the light microscopy facility at Max Planck Florida Institute, and imaged at room temperature using the Zeiss LSM 780 confocal microscope. Z-stacks were acquired using Zen software (2015, 64 bit) and maximum intensity projections were obtained. Dendritic arbors were obtained and quantified using Sholl analysis in Fiji (Image J, NIH). Spine analysis was performed on secondary branches from basal and apical dendrites on the maximum intensity projection images, using a custom MATLAB script. Values were obtained for total, mushroom, thin, and stubby spine density per 100 microns.

### Patch Clamp Electrophysiology

Whole-cell patch-clamp recordings were performed as previously described [11]. Briefly, primary hippocampal neurons at DIV 9-12 were patched using Axon Multiclamp 700 b amplifier, 1440A Digidata digitizer and pClamp software (Axon Instruments, Foster City, CA). Current and voltage recordings were made at 50 kHz, and subsequently filtered at 5 kHz. The recordings were conducted blindly with four groups: untransfected, scrambled LNA Gapmer, ADEPTR Gapmer1, and ADEPTR Gapmer2. Spontaneous excitatory post-synaptic potential currents (sEPSCs) were recorded before and after the bath application of Forskolin under voltage clamp. The current clamp was recorded only to identify the health of the neurons. Only neurons with resting membrane potential of more than −40 mV were used for our analysis.

### Organotypic hippocampal slice cultures and transfection

Organotypic hippocampal slices were prepared from wildtype postnatal 4–7 day old mouse pups of both sexes as previously described [61]. In brief, the animals were anaesthetized with isoflurane, after which the animal was quickly decapitated and the brain removed. The hippocampi were dissected and cut into 350 µm thick coronal hippocampal slices using a McIlwain tissue chopper (Ted Pella, Inc) and plated on hydrophilic PTFE membranes (Millicell, Millipore) fed by culture medium containing MEM medium (Life Technologies), horse serum (20%), L-Glutamine (1 mM), CaCl_2_ (1 mM), MgSO_4_ (2 mM), D-Glucose (12.9 mM), NaHCO_3_ (5.2 mM), Hepes (30 mM), Ascorbic Acid (0.075%), Insulin (1 µg/ml). Slices were incubated at 37 °C in 5% CO_2_. After 7–12 days in vitro, CA1 pyramidal neurons were transfected with biolistic gene transfer using 1.6 µm gold beads (8 mg) coated with plasmids containing 50 µg of total cDNA of interest [62].

### Two-photon fluorescence microscopy and two-photon glutamate uncaging

MNI-caged glutamate uncaging and time-lapse structural imaging of spines were performed using a custom-built two-photon (2p) laser microscope as previously described [63]. 2p-imaging and uncaging was performed using two Ti-sapphire lasers (Coherent, Cameleon) at wavelengths of 920 nm (1.45–1.55 mW under the objective) for imaging and 720 nm (2.7–3.0 mW under the objective) for uncaging. Green fluorescence emission was collected using an immersion objective (LUMPlan FL N 60×, numerical aperture 1.0, Olympus), reflected by a dichroic mirror (565 nm LP) and passed a filter (Chroma, 510nm/70-2p) before entering the fast photoelectron multiplier tubes (PMT) (H7422-40p; Hamamatsu). Fluorescence images were acquired and quantified using TimeHarp 260 Pico card (PicoQuant, Inc) and a custom-built software, FLIMage (Ver 2.0.0) written with #C (https://github.com/ryoheiyasuda/FLIMage_public). Image acquisitions for monitoring long-term structural LTP volume changes were collected by 128 × 128 pixels as a z stack of five frames with 1 µm distance in each frame and averaging 6 scans per frame (frame rate 0.65 Hz). Fast-rate simultaneous image acquisitions with uncaging were collected by 64 × 64 pixels at a single z plane without averaging per frame (frame rate 7.8 Hz). MNI-caged L-glutamate (4-methoxy-7-nitroindolinyl-caged L-glutamate, Tocris) was uncaged with a train of 8–10 ms laser pulses (under the objective, 30 times at 0.5 Hz) near a spine of interest. Experiments were performed at room temperature (24–26 °C) in ACSF solution containing: NaCl (127 mM), KCl (2.5 mM), NaHCO_3_ (25 mM), NaH_2_PO_4_ (1.25 mM), CaCl_2_ (4 mM), glucose (25 mM), tetrodotoxin (1 µM), and 4-MNI-caged L-glutamate (4 mM), bubbled with 95% O_2_ and 5% CO_2_. We examined secondary or tertiary branches of apical dendrites of CA1 pyramidal neurons (located in stratum radiatum) in organotypic cultured hippocampus slices at 11–17 days in vitro.

### Statistical analysis for glutamate uncaging experiments

Spine volume change was calculated by F/F0, in which F0 is the average spine intensity before stimulation. All values are presented as mean ± SEM unless otherwise noted. Number of independent measurements (n[spines/neurons]). One-Way ANOVA followed by Dunnett’s Test was used to compare grouped data sets (Prism 8.3, GraphPad). Data were excluded if signs of poor cellular health or procedural artifacts (for example, dendritic blebbing, displacement of dendrites) were apparent.

### Native and Crosslinked RNA Antisense Purification

RNA antisense purification was performed as previously described [53] with a few modifications. The day before experiments, Streptavidin beads (NEB) were washed twice and blocked in pulldown buffer (Tris-HCl 50mM pH 7.4, NaCl 150mM, EDTA 0.5M, NP-40 0.25%, 1%BSA, DTT 1mM, Superase RNAse inhibitor, Protease inhibitor (Roche) and Phosphatase inhibitor cocktails (Roche)) supplemented with 4% BSA, 20 uM glycogen and 20 uM yeast RNA (ThermoFisher) overnight. On the day of experiments, DIV 15-18 primary hippocampal neurons were treated with Forskolin (50uM) for 30 minutes to boost the endogenous expression of ADEPTR then lysed in pulldown buffer. Cells were scraped and collected into Eppendorf tubes, rotated for 20 min at 4°C, and centrifuged at 1200xg at 4°C for 20 min to pellet the debris. The supernatant was then collected and divided into 400uL reactions. Biotinylated RNA probes were prepared just as in RNA FISH, using Biotin RNA Labeling Mix (Sigma Aldrich). Prior to experiments, probes were heated at 72°C, then kept once ice until added to reaction, and rotated at room temperature for 3 hours. The blocked streptavidin beads were washed an additional time and added to the reaction for an additional hour. After 5 stringency washes, the beads were resuspended with Trizol for RNA extraction or laemmli buffer for protein extraction.

For crosslinked RNA antisense purification, neurons were washed in PBS and crosslinked with 4% PFA for 15 minutes, followed by another PBS wash and finally lysis buffer. Cells were scraped and pooled into one falcon tube for sonication (15 seconds on 15 seconds off for 10 minutes) and gently centrifuged (1200xg, 10 min, 4°C) to clear the debris. Supernatant was then subjected to pulldown as described above. Stringency washing was done with pulldown buffer lacking BSA and an NaCl concentration of 600mM. Beads were then resuspended with Tris buffer containing 10ug/uL RNAse A, incubated at 37°C for 60 min, and pelleted. The supernatant was treated with Superase RNAse Inhibitor and transferred to an ultracentrifuge tube for ultracentrifugation (70K RPM, 4 hours, 4°C) with a 1M sucrose gradient. The supernatant was then decanted and the pellet was suspended in Trizol for total RNA extraction, bioanalyzer measurements and RNA sequencing.

### RNA EMSA

EMSA was performed using the LightShift® Chemiluminescent RNA EMSA Kit (Thermo Scientific) as per instructions. Primary hippocampal neuron lysate was prepared as described in the pulldown assay. 5ug protein was incubated with 100 ng of RNA (3’ end labeled biotinylated probes were synthesized from IDT). The IRE control was used from the kit. Binding reactions were then run on a 6% polyacrylamide TBE gel in 0.5X TBE buffer and transferred onto a nylon membrane for streptavidin-HRP-based detection of bands.

### Silver staining and Proteomics

Proteins were separated by SDS-PAGE, washed in water, and fixed overnight (10% glacial acetic acid, 30% Ethanol) on an end-to-end shaker. Total protein bands were visualized using Pierce Silver Staining Kit (Thermofisher). For mass spectrometry, a large scale pulldown from 3 combined experiments was run on SDS page, lanes corresponding to Antisense and Sense probe groups, were cut out and processed for LC/MS-MS. Unique peptides were ascertained using Scaffold software.

### Western Blotting

Protein was isolated from whole cells, cell fractions, or RNA pulldown assays as described above and ultimately prepared in laemmli buffer with 10% B-mercaptoethanol and 1% SDS. Equal volumes of protein were loaded for SDS-PAGE followed by overnight transfer to methanol-activated PVDF membranes. Membranes were then probed for Ankyrin-B (Santa Cruz), Spectrin-a (Santa Cruz), Actin (Abcam), GAPDH (Abcam). The target proteins were detected using anti-rabbit or anti-mouse secondary antibodies (Cell Signaling) at 1:5000 dilution, then visualized by Pierce™ ECL Western Blotting Substrate (Thermofisher). Blots were quantified using FIJI ImageJ.

### Lentivirus preparation

PLL3.7 shRNA plasmid with lentivirus backbone was kindly provided by the Xu lab. ADEPTR shRNA (mature sequence: 5’GCTAGGTGCAAGATGGAA3’) was then cloned into this backbone, replacing the scrambled sequence, and verified by Sanger Sequencing (Genewiz). Each plasmid was co-transfected with helper plasmids (pMDL, pRev, pVSVg) into HEK293T cells using Mirus reagent. The media was collected 48-72 hours later. Viral particles were concentrated and ultra-centrifuged (25K RPM, 2.5 hours, 4°C) with sucrose gradient. The viral pellet was then re-suspended in sterile PBS and stored at −80°C. Viral titer was determined by measuring GFP fluorescence after infection of HEK 293 cells with serial dilutions of the virus.

### Statistical Analysis

Statistical analysis was performed in R and Prism 8. Data are represented by the mean and error bars represent SEM. Statistical tests performed are Unpaired Two-tailed T-test, One-Way, Two-Way ANOVA with Tukey’s HSD, Dunnett’s Test, Holm’s correction post hoc test, or pairwise-T-test, as indicated. N represents the number of independent samples for each experiment, unless stated otherwise.

### Data Availability

RNAseq data related to Figure S1 were deposited to NCBI Gene Expression Omnibus with the accession number GSE157388, and all relevant data are available from the authors upon reasonable request.

## Acknowledgments

We would like to acknowledge the help of Drs. Naomi Kamasawa and Long Yan from Max Planck Florida Institute for help with Electron Microscopy and Fluorescence Microscopy, respectively. We also acknowledge Dr. Miklos Kepiro for help with image analysis in ImageJ, Dr. Pabalu Karunadharma at Scripps Florida Genomics Core for preparation of libraries for RNA sequencing, and the Xu lab at Scripps Florida for kindly providing the PLL3.7 shRNA plasmid. Finally, we gratefully acknowledge the funding support from NIH (5R01MH094607-05, 1R21DA039417-01A1 and 1R01MH119541-01A1) to carry out this work.

## Supplementary Materials

Figures S1-S14

Supplementary Tables S1-S12

## Supplementary Figures

**Supplementary Figure 1:**
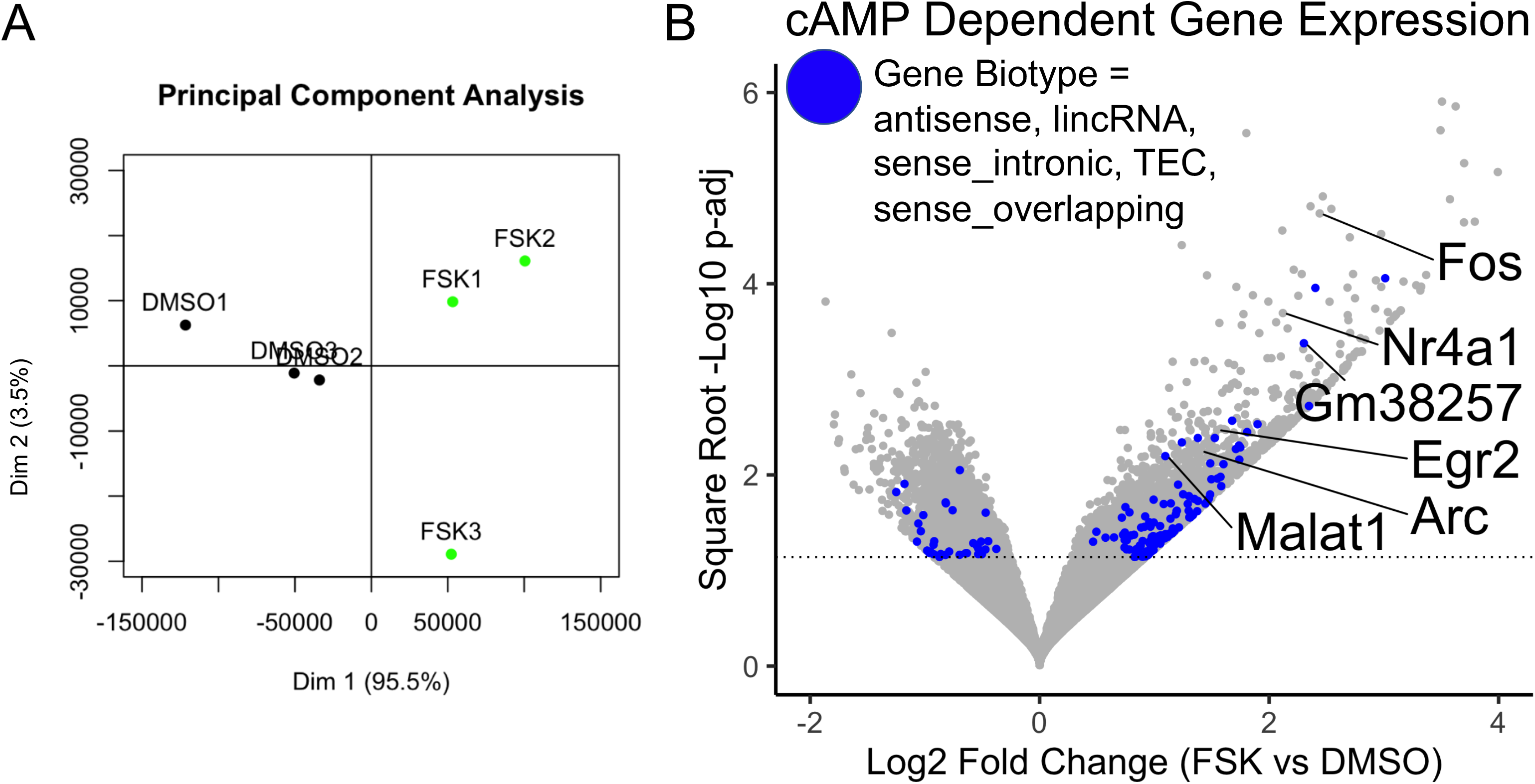
RNASeq Analysis of Hippocampal Neurons. **(A)** Principal Component Analysis of RNAseq data from Forskolin or vehicle treated DIV 16 hippocampal neurons. **(B)** Volcano plot indicating DEseq-identified significant genes. Gene biotypes were obtained using Ensembl BiomaRt R package. Positive markers for cAMP-dependent gene expression (Fos, Egr2, Nr4a1, and Arc) are indicated. N=3 biological replicates per group.

**Supplementary Figure 2:**
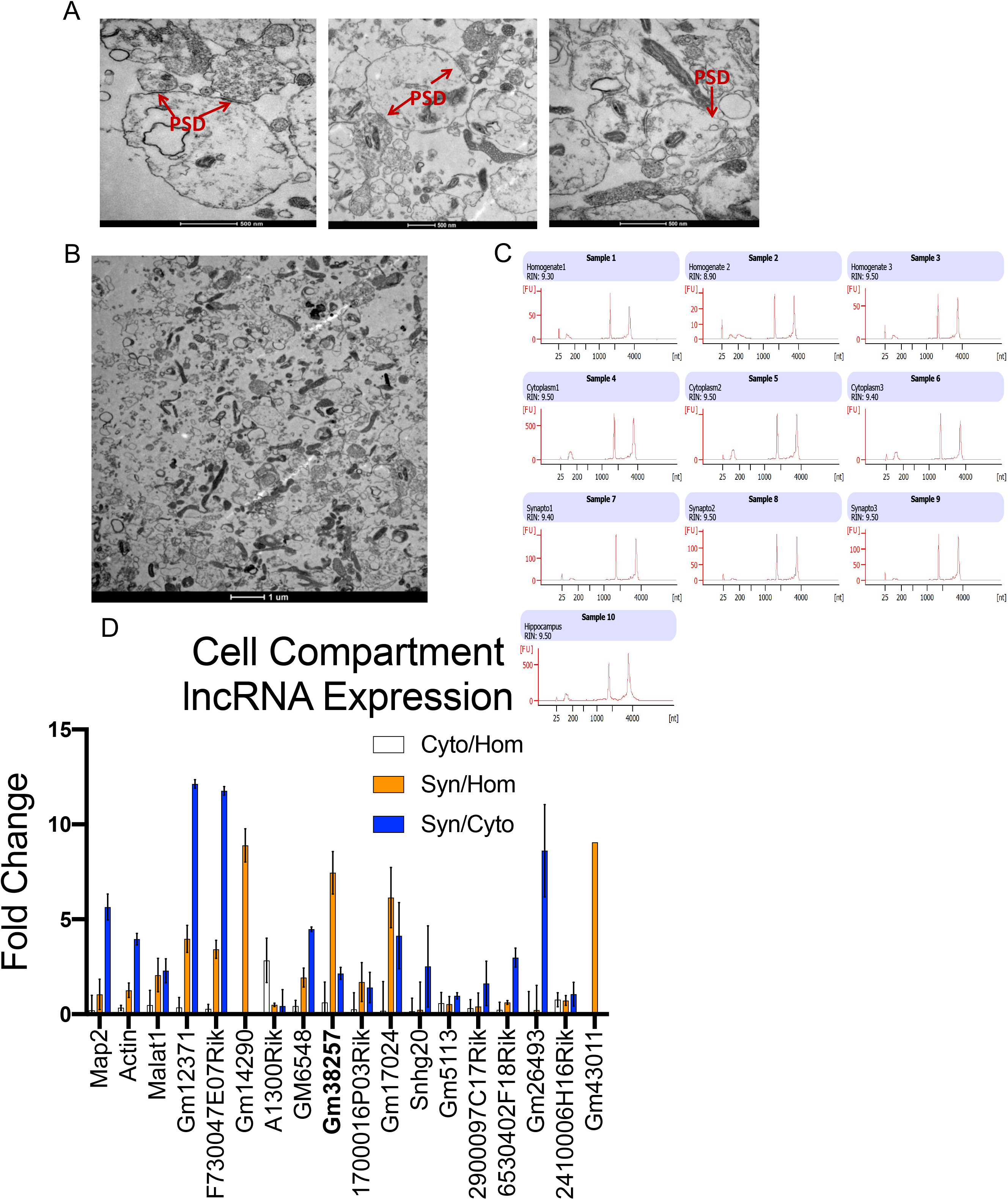
Quality control of synaptoneurosome preparation, RNA extraction, and lncRNA screening. **(A)** Three transmission electron micrographs of synaptoneurosome preparations with post synaptic densities (PSD) indicated. **(B)** Zoomed out view of transmission electron micrograph of synaptoneurosome preparation showing no presence of nuclei or intact cells. **(C)** Bioanalyzer runs indicating RNA quality from hippocampal neuron homogenate, cytoplasm, and synaptoneurosome preparations. Total hippocampus was used as a positive indicator of RNA integrity. **(D)** qRT-PCR screen for lncRNA enrichment in synaptoneurosomes, as compared to cytoplasm or homogenate. N=3 per group except for Gm43011 (N=1). Error bars represent SEM.

**Supplementary Figure 3:**
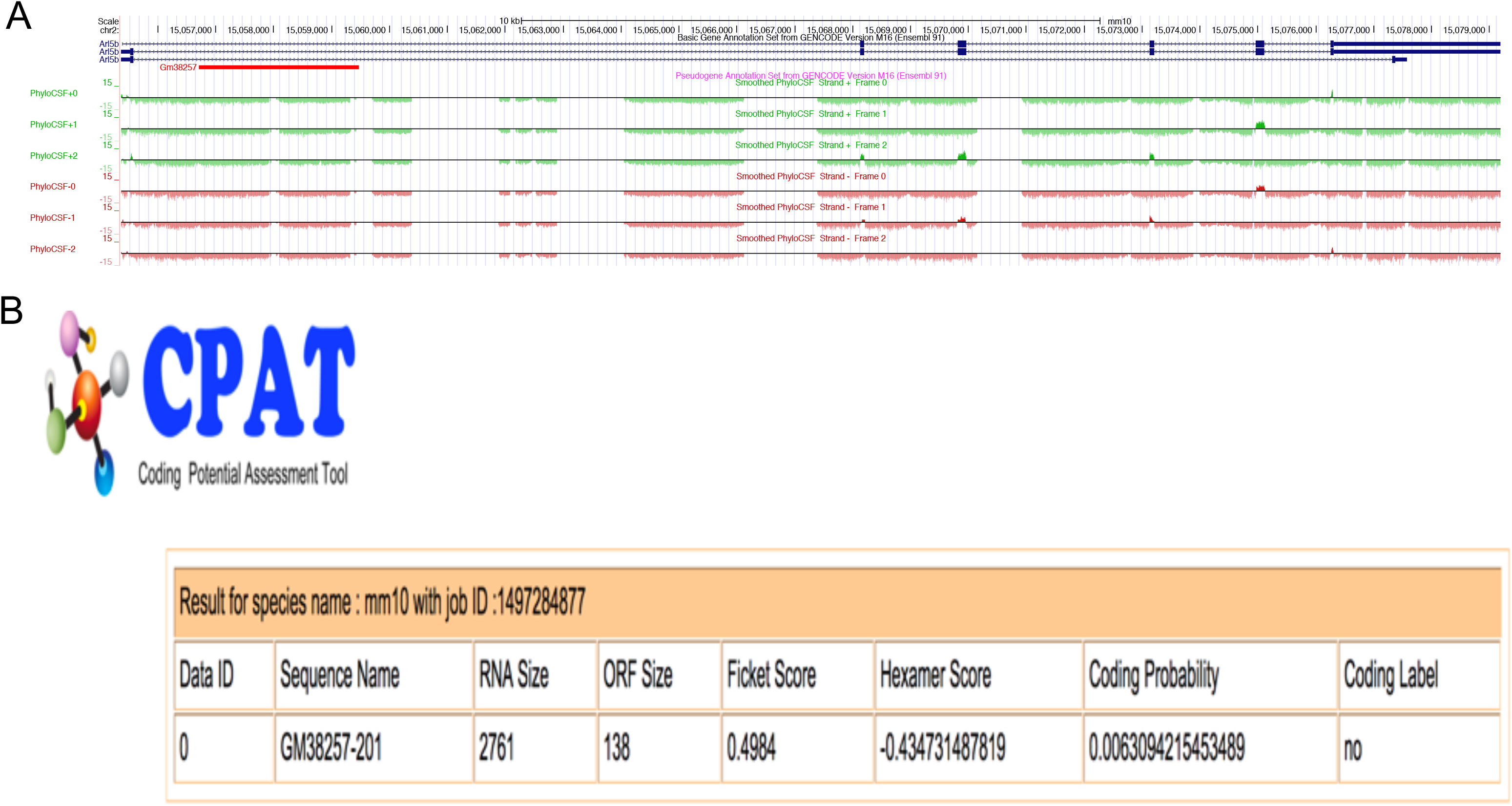
Gm38257 is an intronic localized lncRNA within Arl5b and as no protein coding potential. **(A)** UCSC genome browser mm10 track of Arl5b intron with PhyloCSF 6-frame translation indicating no predicted open reading frame within Gm38257. **(B)** Coding Potential Assessment Tool indicating one ORF within Gm38257 that has a no-coding label.

**Supplementary Figure 4:**
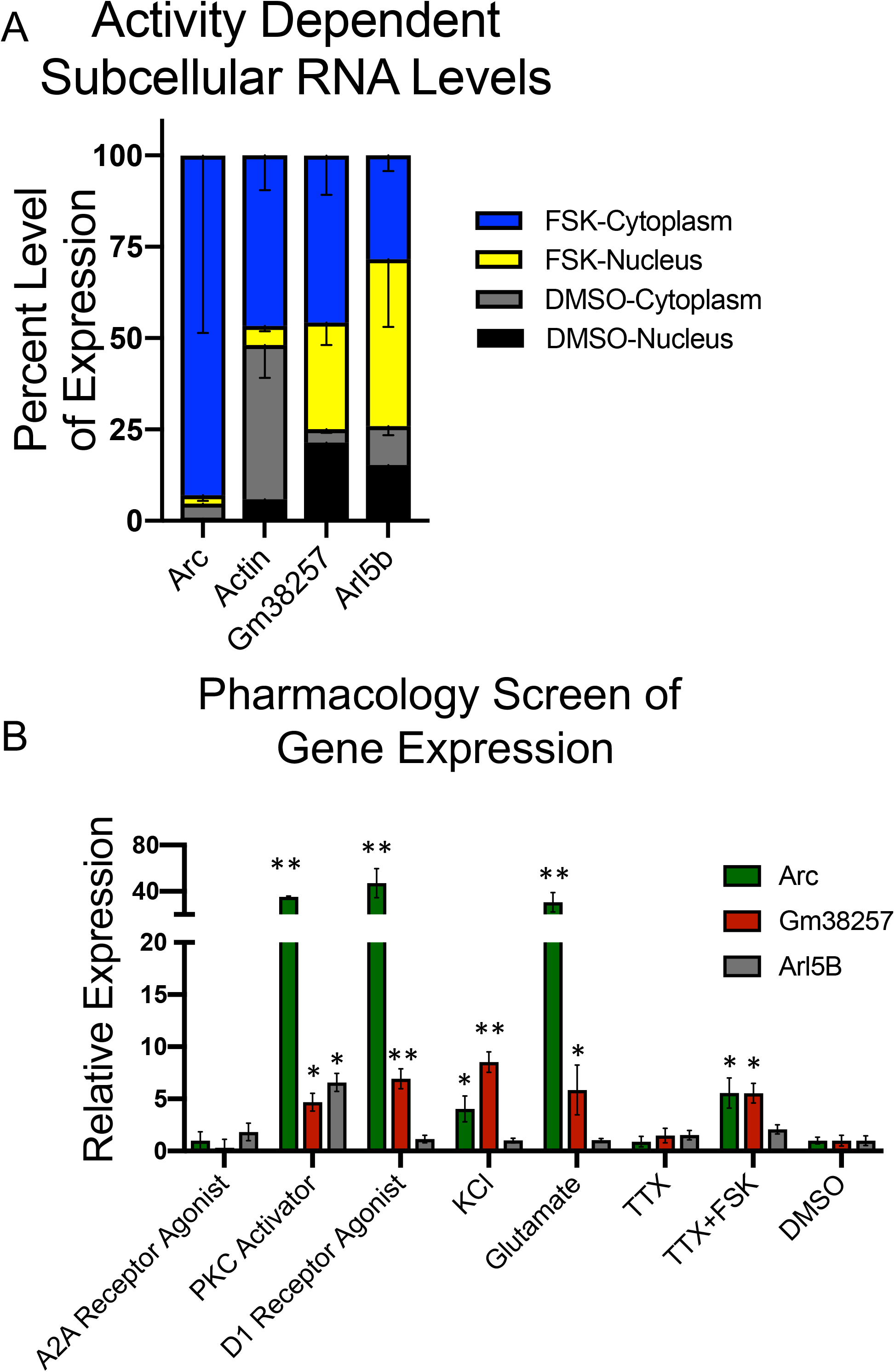
cAMP-dependent ADEPTR is dendritically localized as a nascent RNA and is expressed independently of Arl5b. **(A)** Nuclear/cytoplasmic isolation of DMSO or FSK treated neurons followed by RNA extraction and qRT-PCR (N=3) per group. **(B)** DIV 15-18 primary hippocampal neurons were treated with indicated pharmacological agents for 30 minutes prior to RNA extraction and qRT-PCR analysis. Arc was used as a positive control for gene expression. N=3-4 per group. *p<0.05, **p<0.01. Student’s T-test compared to DMSO control.

**Supplementary Figure 5:**
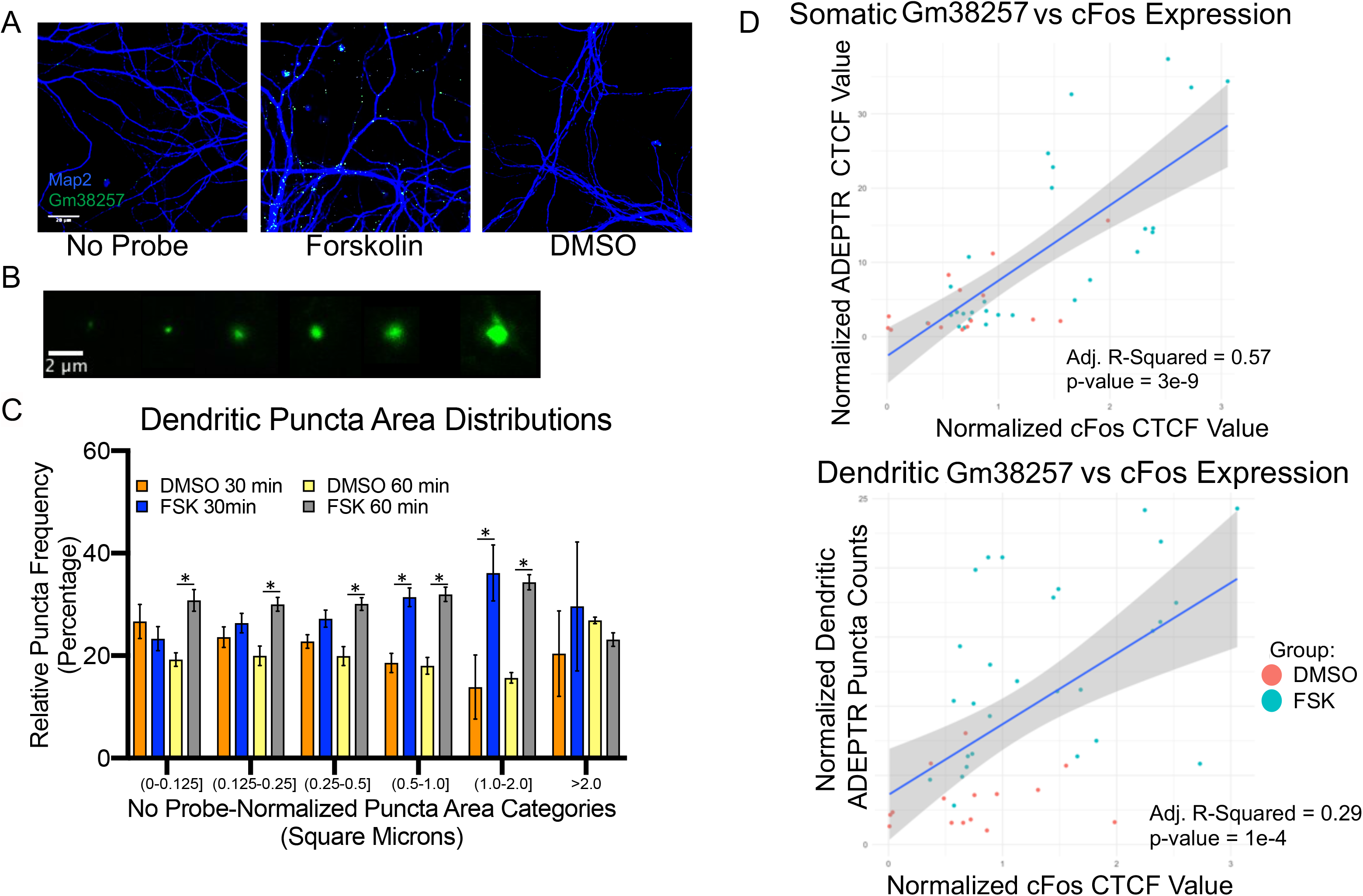
ADEPTR is expressed in distal neuron processes in large sized puncta a cAMP/PKA dependent manner. **(A)** Confocal images of ADEPTR FISH on Map2 labeled distal processes from hippocampal neurons. No probe was used as a negative control. Scale bar is 20 microns **(B)** Representative insets of ADEPTR puncta ranging from 0.1 to 2 square microns. **(C)** Extra somatic puncta were acquired and by a custom ImageJ macros script; the areas of each individual punctate were sorted into discrete categories, averaged for each neuron, and plotted as a percentage of all puncta in that category. N=17-24 per group from 3 independent experiments. **(D)** Regression analysis of cFos nuclear fluorescence vs (top) Gm38257 somatic fluorescence and (bottom) ADEPTR dendritic puncta counts from Figure 1. *p<0.05, **p<0.01. Student’s T-test. Error bars for all graphs represent SEM.

**Supplementary Figure 6:**
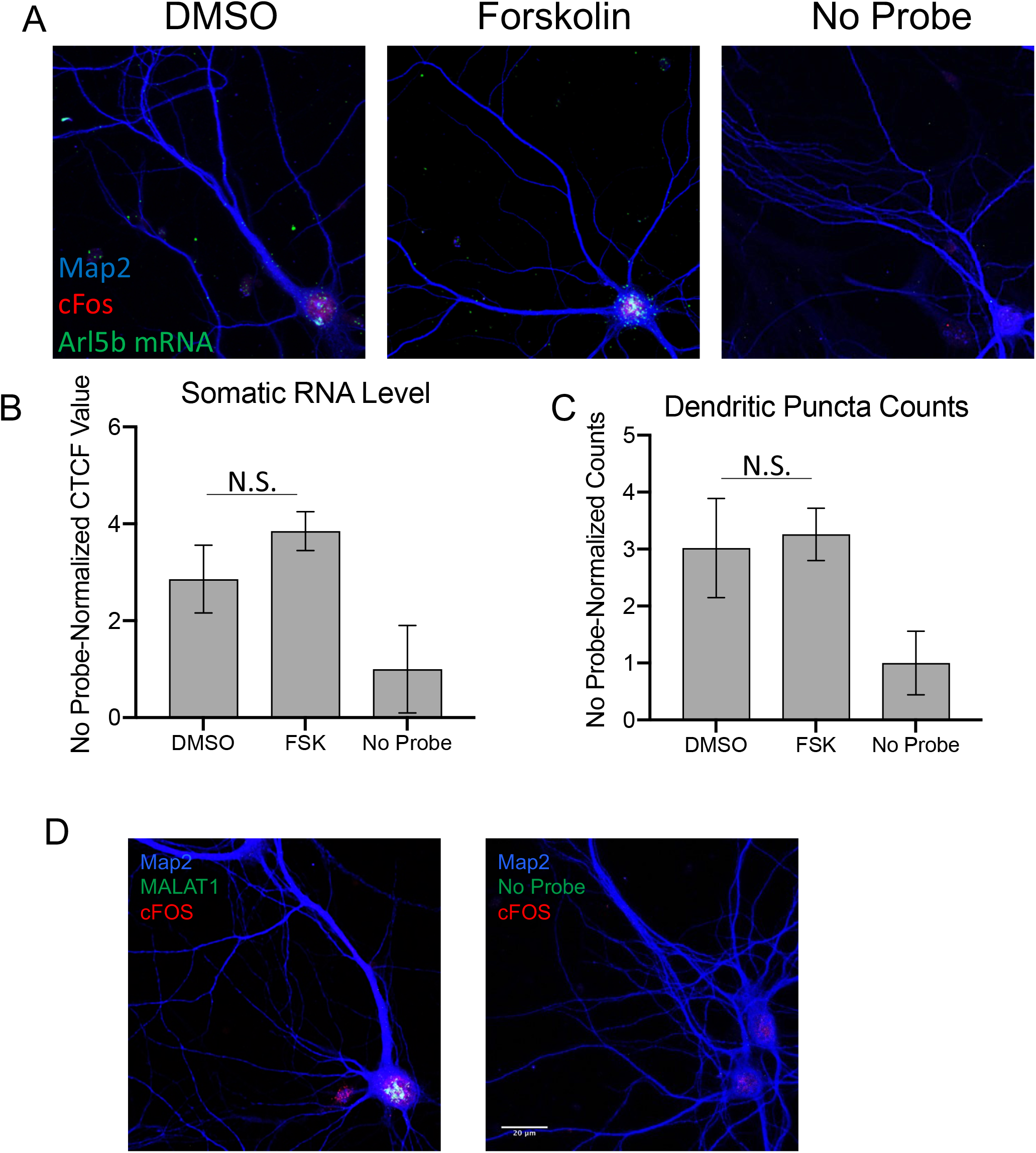
Arl5b mRNA is not dendritically localized in a cAMP-dependent manner. **(A)** Primary hippocampal neurons were processed for FISH as in Figure 1 but treated with Arl5b antisense DIG-labeled riboprobes. **(B)** Quantification of somatic Arl5b fluorescence and **(C)** dendritic RNA puncta counts as in Figure 1. N=5-7 per group. **(D)** Malat1 and No probe FISH experiments were included as technical controls. One Way-Anova, Tukey’s HSD. Error bars represent SEM. Scale bar is 20 microns.

**Supplementary Figure 7:**
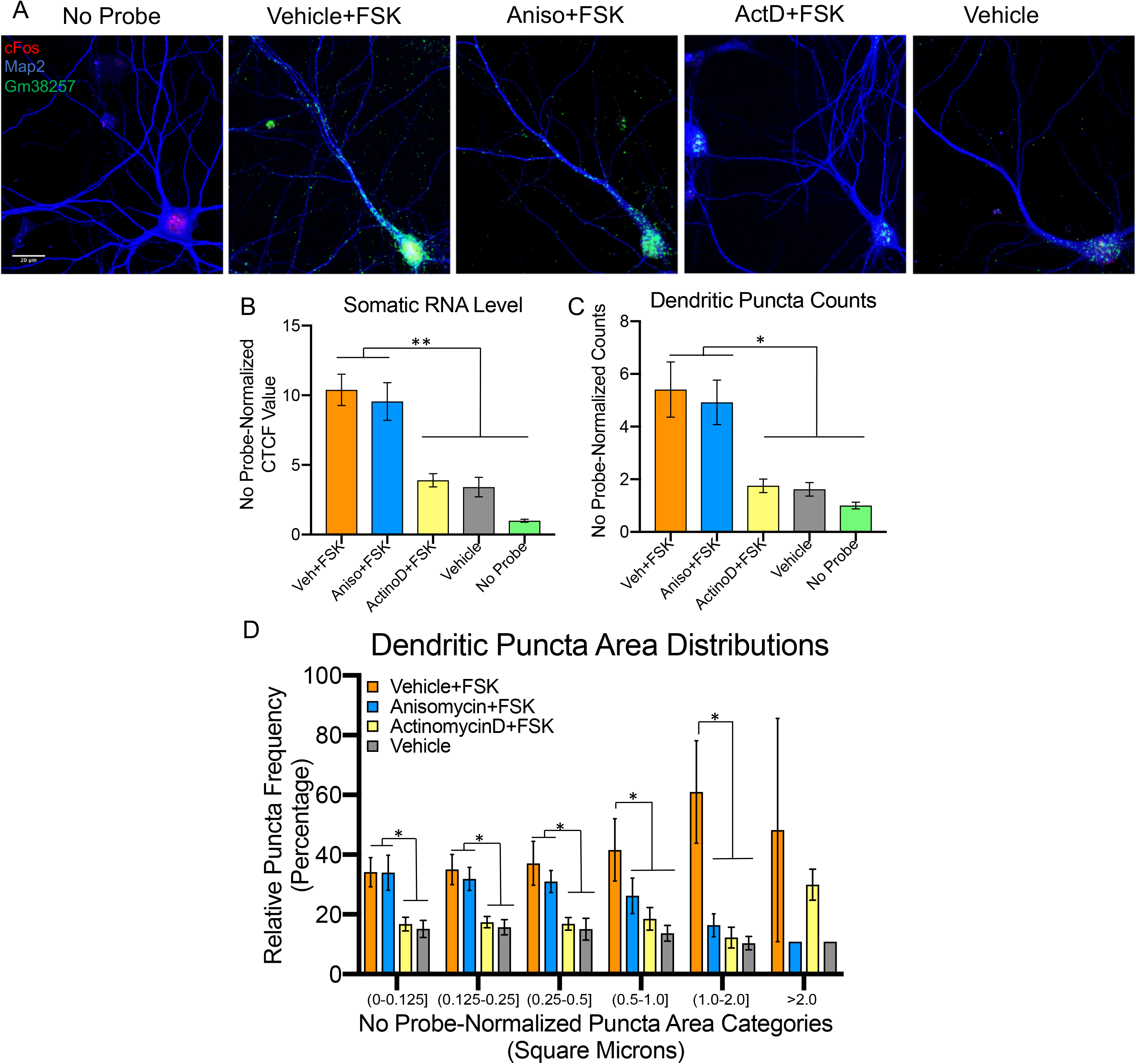
cAMP-dependent dendritic localization of ADEPTR is dependent on transcriptional but not translational activity. **(A)** Primary hippocampal neurons were treated with vehicle, Anisomycin, or ActinomycinD for 30 minutes, followed by Forskolin for another 30 minutes, prior to FISH. Vehicle (DMSO) served as pharmacological control. **(B)** Quantification of somatic ADEPTR levels, **(C)** dendritic ADEPTR puncta, and **(D)** puncta area distributions. N=8-15 from 2 independent experiments. *p<0.05, **p<0.01. Pairwise T-test with Holm correction. Error bars represent SEM.

**Supplementary Figure 8:**
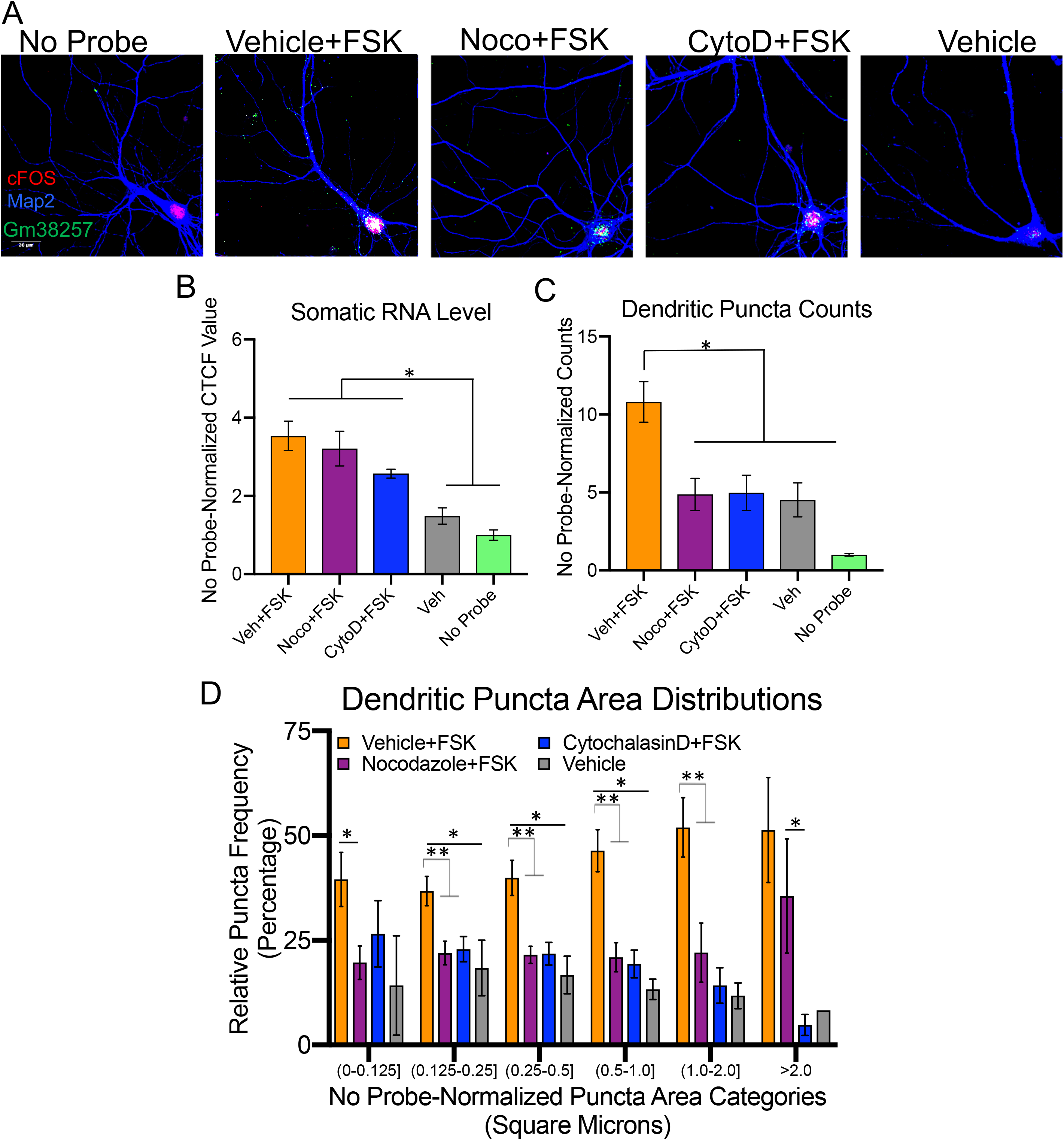
cAMP-dependent dendritic localization of ADEPTR is dependent on microtubule and actin polymerization. **(A)** Primary hippocampal neurons were treated with vehicle, Nocodazole, or CytochalasinD for 30 minutes, followed by Forskolin for another 30 minutes, prior to FISH. Vehiclen(DMSO) served as pharmacological control. **(B)** Quantification of somatic ADEPTR levels, **(C)** dendritic ADEPTR puncta, and **(D)** puncta area distributions. N=5(DMSO), 7(Vehicle), 20(Veh+FSK), 16(Noco+FSK), 13(CytoD+FSK) from 2 independent experiments. *p<0.05, **p<0.01. Pairwise T-test with Holm correction. Error bars represent SEM. Error bars represent SEM.

**Supplementary Figure 9:**
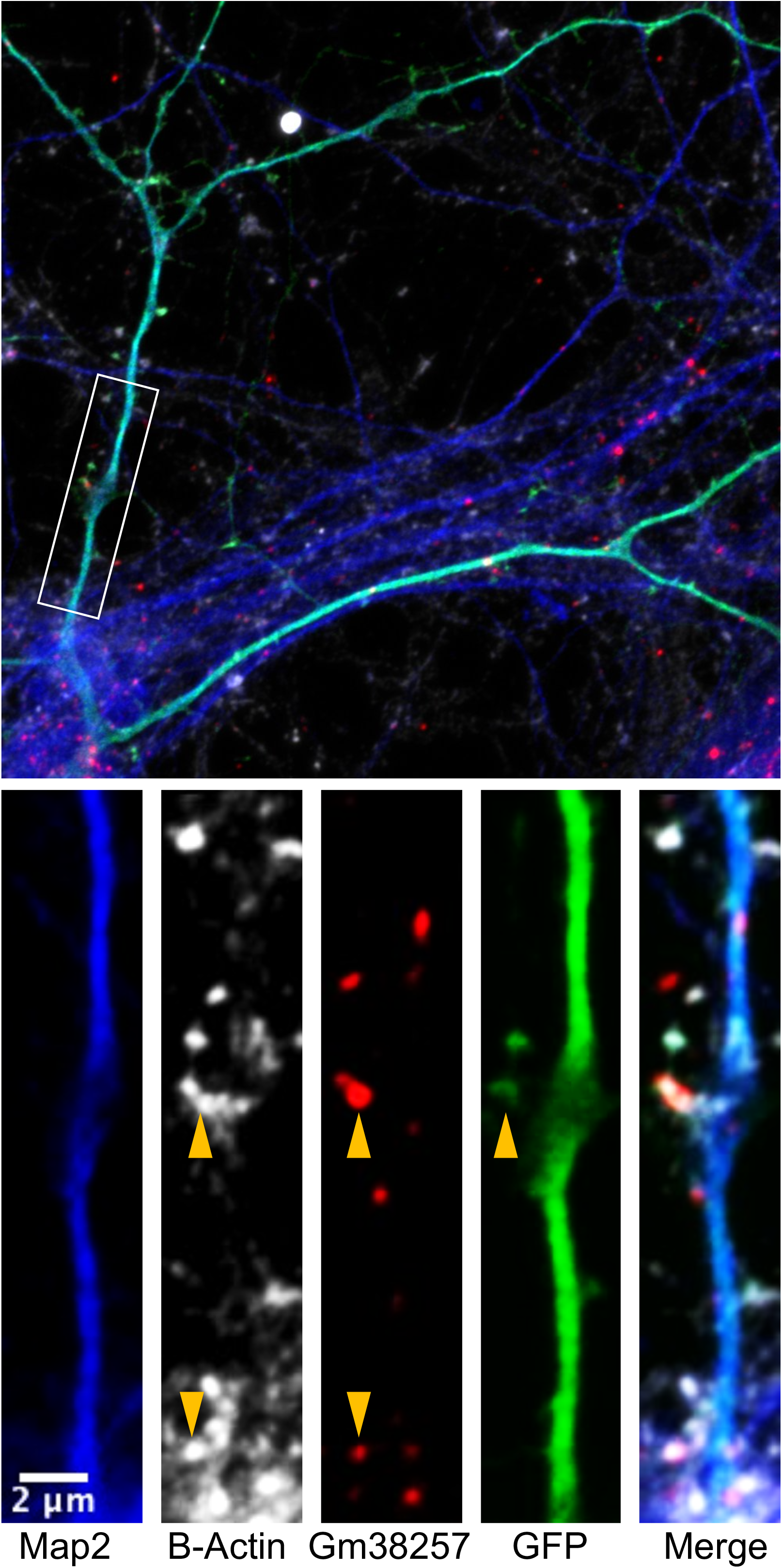
ADEPTR expression is observed within dendritic spines. Inset shows super Resolution image of GFP labeled Forskolin treated neuron labeled with Map2 (blue), B-actin (white), and lncRNA Gm38257 (Red).

**Supplementary Figure 10:**
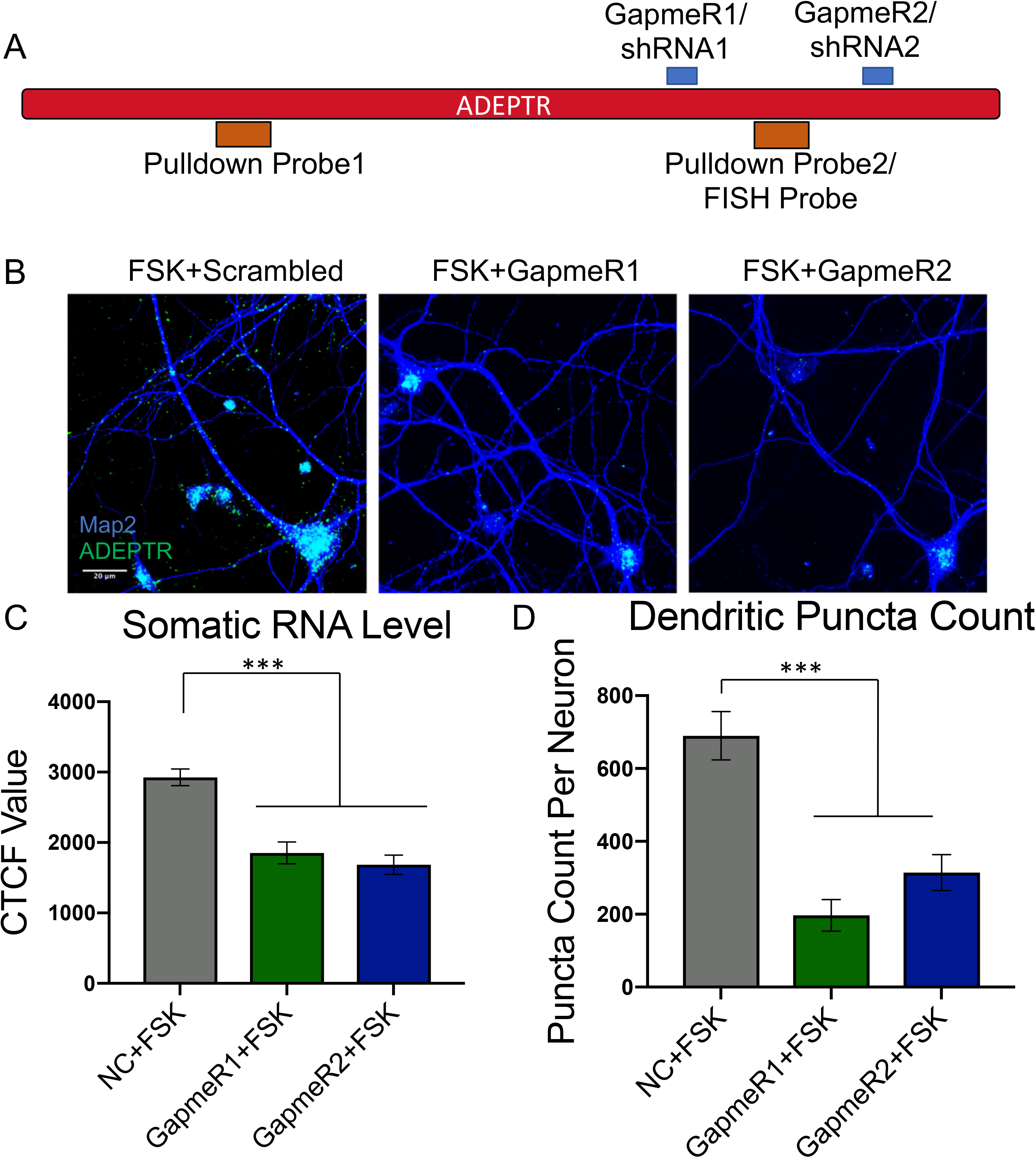
ADEPTR knockdown by two independent LNA GapmeRs. **(A)** Schematic of positions of GapmeRs, shRNAs, and probes for ADEPTR. **(B)** Neurons were treated with GapmeRs 48 hours prior to Forskolin stimulation, followed by FISH. **(C)** Quantification of somatic ADEPTR levels and **(D)** dendritic ADEPTR puncta numbers. N=14-16 neurons from two independent experiments. ***p<0.001 One-Way ANOVA, Tukey’s HSD. Error bars represent SEM. Scale bar is 20 microns.

**Supplementary Figure 11:**
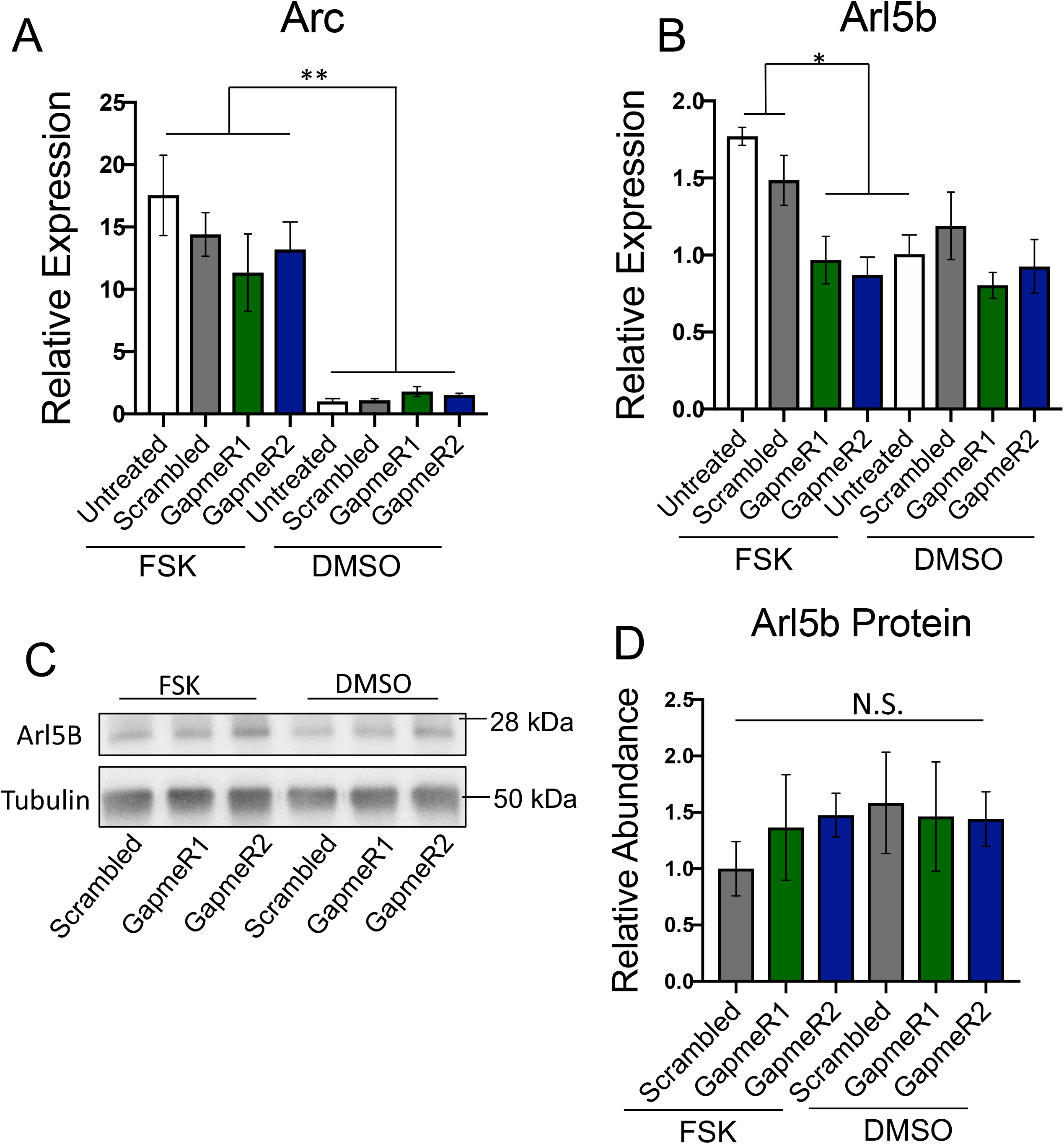
ADEPTR deficiency does not impact Arc mRNA or Arl5b protein levels. **(A)** qRT-PCR analysis of Arc and **(B)** Arl5b expression in Forskolin or DMSO-treated hippocampal neurons from same samples as in Figure 2B. N=4-6 per group. **(C)** Western blot analysis of total protein from hippocampal neurons treated as in Figure 2. **(D)** Quantification of Arl5b protein levels, N=3. * p<0.05, **p<0.01, One Way-ANOVA, Tukey’s HSD.

**Supplementary Figure 12:**
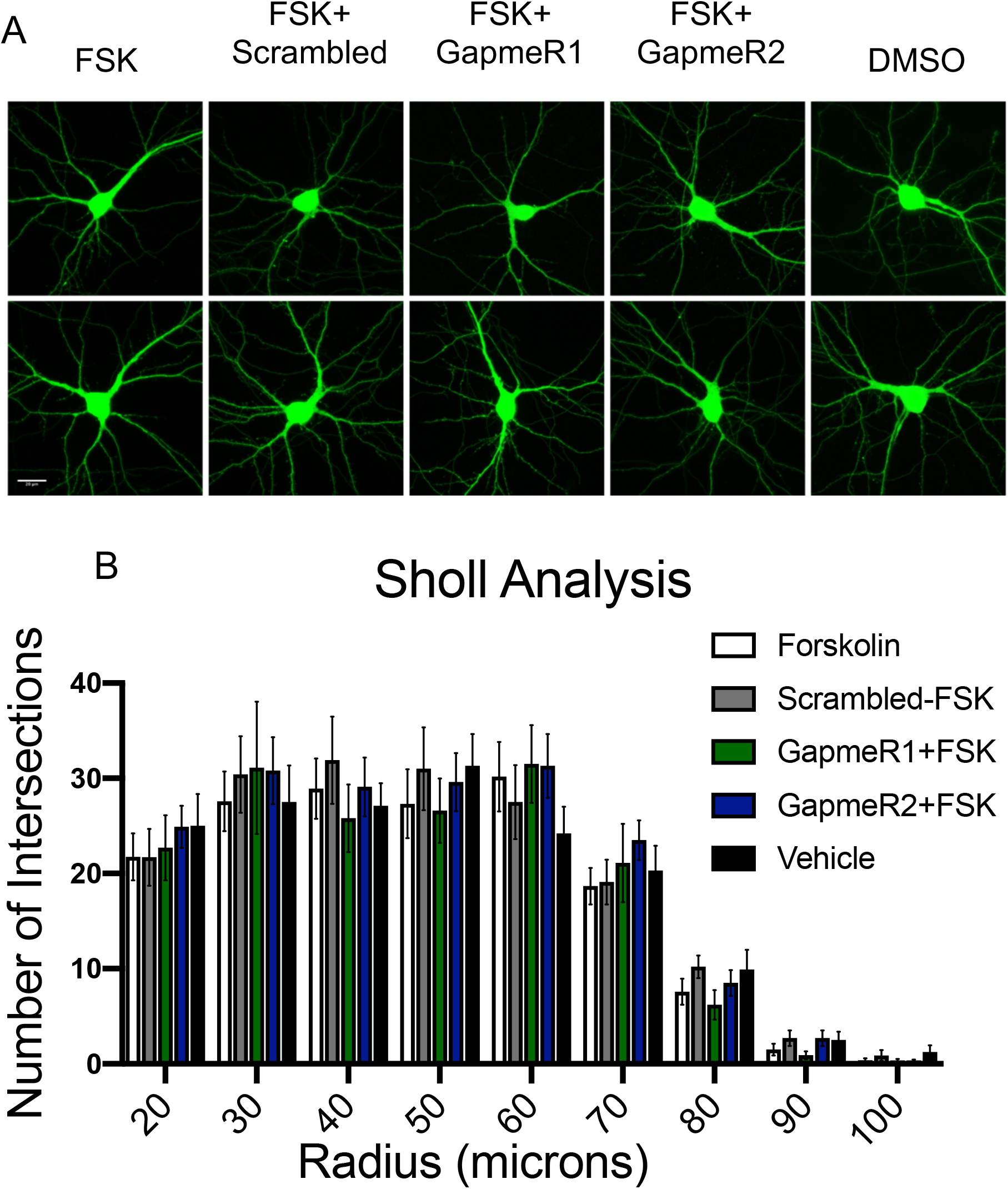
ADEPTR deficiency does not influence neuronal arborization. **(A)** Live imaging from GFP labeled neurons with the indicated treatments. **(B)** Sholl analysis was performed using Fiji ImageJ plugin by counting the number of intersections from the soma beginning at 20 microns. N=10-12 neurons per group. One-way ANOVA. Error bars represent SEM.

**Supplementary Figure 13:**
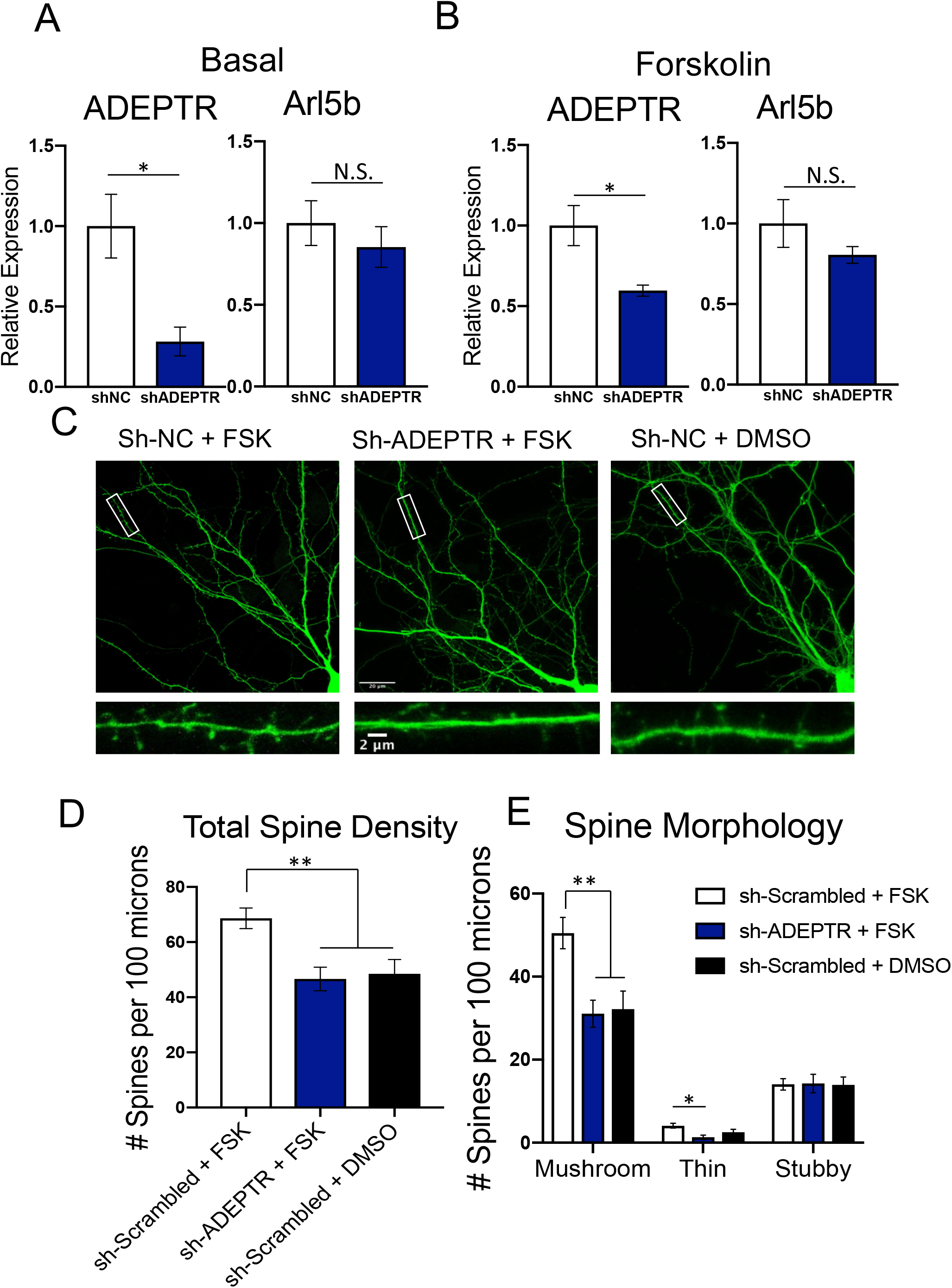
ADEPTR deficiency by shRNA suppresses cAMP-dependent changes in dendritic spine density and morphology. **(A)** An shRNA cassette targeting the same region as GapmeR2 was cloned into PLL3.7 vector with a GFP reporter, followed by packaging into lentiviral vectors. Neurons were transduced with virus at DIV 3 and RNA was collected at DIV 14. Graphs indicate qPCR validation of ADEPTR (left) and Arl5b (right) in Basal and **(B)** Forskolin conditions. N=6-7 per group from 2 independent experiments. **(C)** Primary hippocampal neurons were sparsely labeled by transfecting shRNA plasmids at DIV 14 for 72 hours, treated with Forskolin or DMSO, then subjected to live imaging by Zeiss 780 confocal microscope. **(D)** Analysis of spine density and **(E)** morphology as performed by MATLAB custom script. N=19-31. *p<0.05, **p<0.01. One Way-ANOVA, Tukey’s HSD. Error bars represent SEM.

**Supplementary Figure 14:**
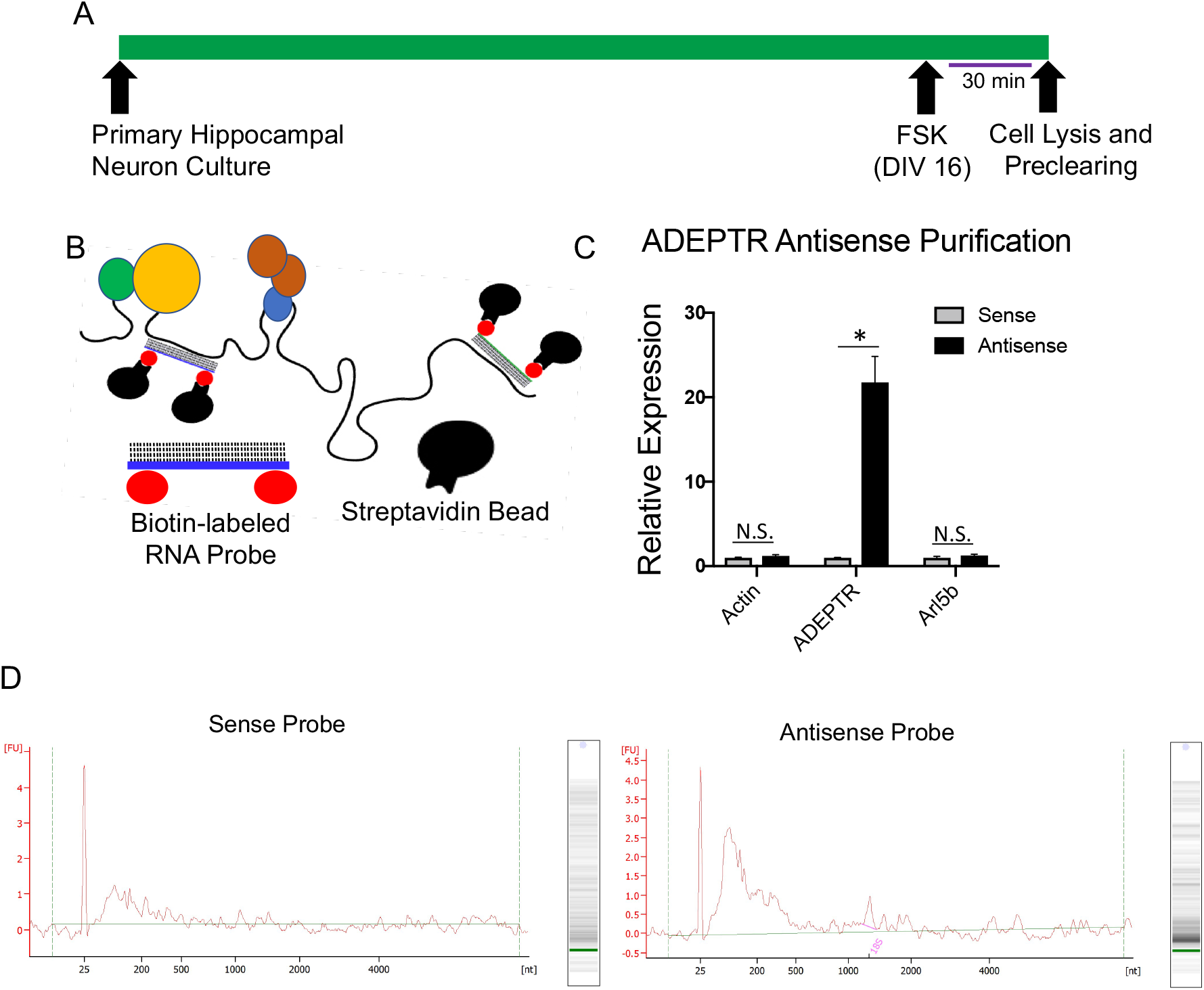
ADEPTR native and crosslinked pulldown validation. **(A)** Schematic demonstrating the time course for antisense purification of ADEPTR. **(B)** Schematic of antisense purification strategy; 300-400 nt probes corresponding to ADEPTR were cloned from cDNA into a PCRII TOPO vector for sense and antisense in vitro transcription using nucleotides containing biotin-labeled uracil. Target RNA was isolated by biotin capture with streptavidin magnetic beads, followed by Trizol RNA isolation. **(C)** Expression profile of corresponding gene in antisense and sense probe groups. Exogenous firefly luciferase mRNA was spiked into the preparation for normalization during relative quantification qRT-PCR. N=5-8 reactions per group. * p <0.05, Student’s T-Test. **(D)** Bioanalyzer (Pico kit) results of the crosslinked ADEPTR antisense purification with sense control (left) and antisense probe (right)

